# A CRISPR-Cas9 platform for primary human hepatocytes enables arrayed screening and *in vivo* validation of HBV host factors

**DOI:** 10.64898/2026.08.10.743996

**Authors:** Ansgar F. Stenzel, Antonis Athanasiadis, Georgios Dangas, Paul Park, Anthi Maslarinou, Evgenia Moschogianni, Allan Henrique Depieri Cataneo, Catherine A. Freije, Yichen Zhou, Kenneth C. Levenson, Corrine Quirk, Chenhui Zou, William M. Schneider, Adriano Aguzzi, Charles M. Rice, Ype P. de Jong, Eleftherios Michailidis

## Abstract

More than two million deaths annually are attributed to liver-related conditions, making primary human hepatocytes (PHH) an invaluable *in vitro* model for studying liver pathophysiology and the molecular mechanisms underlying hepatic diseases. However, because PHH do not proliferate in culture, CRISPR gene editing has been highly inefficient. Here, we report lipofection- and lentivirus-mediated protocols for CRISPR-Cas9 delivery in mouse-passaged primary human hepatocytes (mpPHH), a system that enables PHH expansion in liver-humanized mice. We achieve robust gene editing efficiencies exceeding 90% in mpPHH while maintaining cell viability. We demonstrate the utility of these protocols by disrupting *CYP3A4* to impair xenobiotic metabolism and by showing that edited mpPHH efficiently engraft and expand in mice, generating liver-humanized animals. We establish the feasibility of arrayed CRISPR screening in mpPHH using an 85-gene screen to identify host factors influencing hepatitis B virus (HBV) infection, and validate key findings in humanized mice by targeting the HBV entry receptor *SLC10A1* (NTCP), which reduced viral infection *in vivo*. Our methodology enables scalable genetic manipulation of mpPHH, opening new avenues for HBV research and liver disease modeling.

## Introduction

Hepatocytes are the parenchymal cells of the liver, serving pivotal physiological functions in metabolism, storage, lipid and protein secretion, and detoxification^1^. Their metabolic functions put them at the center of steatotic liver disease pathogenesis, and their importance in xenobiotic metabolism frequently causes drug-induced liver injury (DILI), a significant challenge in drug development^2^.

Beyond these metabolic roles, hepatocytes are targets of several globally distributed hepatotropic infections, including hepatitis B virus (HBV) and hepatitis C virus (HCV), which can lead to chronic infection and liver cancer^3^. HBV alone chronically infects approximately 254 million people worldwide and causes an estimated 1.1 million deaths annually from cirrhosis and hepatocellular carcinoma^4^. Despite the availability of vaccines and suppressive treatments, functional cure rates remain low, highlighting the urgent need to systematically identify host factors that could serve as novel therapeutic targets. However, our understanding of HBV-host interactions is limited by the lack of scalable genetic approaches in physiologically relevant hepatocyte models.

Advancing our understanding of hepatocyte function and pathophysiology is therefore urgent. However, human cell lines lose hepatic differentiation^5^, and animal models diverge substantially from human hepatocyte biology, particularly in xenobiotic metabolism and susceptibility to hepatotropic pathogens such as HBV. The “gold standard” cell culture system for studying hepatocyte biology is primary human hepatocytes (PHH) isolated from human donors. Yet, PHH are challenging to culture and have several experimental limitations. Foremost among these is that they remain functional in culture for only a limited time (1-2 weeks) and start de-differentiating after plating^6^. Additionally, reproducibility is constrained by limited availability and high donor-to-donor variability. To overcome these restrictions, protocols have been developed to transplant PHH into immunodeficient mice with liver injury, where they can proliferate and repopulate the murine liver parenchyma, thereby forming human hepatocyte chimeric mice^7–11^. Human hepatocytes can then be isolated as so-called mouse-passaged PHH (mpPHH), which form stable long-term (>2 months) cultures^12^. In one such chimeric model using *Fah*^-/-^ NOD *Rag1*^-/-^ *Il2rg*^null^ (FNRG) mice, cryopreserved PHH expand 100-200-fold and can be serially passaged in mice, providing a practically unlimited resource for hepatocyte-related studies^12^.

While both PHH and mpPHH represent valuable models for studying hepatocyte biology, efficient genetic manipulation in these systems remains technically challenging. CRISPR screening has emerged as a powerful technology for unbiased functional genomics, rapidly advancing our understanding of diverse biological processes^13^. In these screening technologies, CRISPR-Cas9 single guide RNA (sgRNA) complexes direct site-specific DNA breaks that are repaired imperfectly by non-homologous end joining, frequently introducing frameshift mutations that disrupt protein function. However, developing CRISPR screening platforms in physiologically relevant hepatocyte models remains challenging. Since PHH do not proliferate in culture, they cannot support conventional pooled CRISPR screens that depend on cellular proliferation for positive or negative selection.

Arrayed CRISPR screening offers an alternative approach that uses parallel, predefined gene perturbations in separate wells^14–16^. This format enables diverse phenotypic readouts, including fluorescence, luminescence, imaging, and proteomics, without requiring DNA sequencing. In addition, arrayed CRISPR screening does not require cell proliferation, making it uniquely suited for non-dividing primary cells such as PHH. While arrayed screens typically cover smaller gene sets than pooled approaches, they enable phenotypic readouts that pooled formats cannot support, including phenotypes masked by paracrine effects between neighboring knockouts and transient phenotypes that resolve before pooled readouts are taken. Therefore, developing an effective arrayed CRISPR screening platform for hepatocytes could significantly advance functional genomics research in liver biology.

Despite mpPHH solving the availability and expansion limitations of traditional PHH, no optimized CRISPR-Cas9 methods exist for genetic manipulation of these cells. A broadly applicable *in vitro* method for CRISPR-Cas9 gene disruption and screening in PHH would unlock the full potential of this physiologically relevant model. However, PHH are non-dividing *in vitro*, so successful CRISPR-Cas9 gene disruption requires balancing two main objectives: as many cells as possible should carry a genotype that leads to protein loss-of-function, while delivery of the Cas9-sgRNA ribonucleoprotein (RNP) complex must have minimal detrimental effects on cell health. Prior efforts to edit PHH have relied primarily on electroporation. Two recent studies demonstrated RNP delivery via electroporation, with editing efficiencies of 50% and 60%, respectively^17,18^, but at substantial cost to cell viability; only approximately two-thirds of PHH survived the electroporation conditions required to achieve this editing^17^. Additionally, neither study demonstrated long-term culture viability or downstream manipulation of edited PHH. Beyond its impact on cell health, electroporation is incompatible with flexible experimental designs and does not scale to high-throughput formats. Viral vector delivery has also been attempted, but reports in cultured PHH have shown limited editing efficiency^19–21^. Lipid-based chemical transfection (lipofection) is another well-established delivery method (reviewed in^22^) that may better preserve cell health while supporting scalable, arrayed formats. We therefore hypothesized that an optimized lipofection-based protocol for CRISPR-Cas9 gene editing in mpPHH could maintain cell viability, enable flexible long-term experimental conditions, and be scaled up for arrayed high-throughput screens.

Here, we establish a scalable CRISPR-Cas9 platform for gene disruption and functional genomics in primary human hepatocytes that supports efficient editing while preserving hepatocyte viability and function. Edited mpPHH can be expanded in humanized mice, enabling phenotyping both *in vitro* and *in vivo*. Applying this platform to arrayed CRISPR screening, we identify host factors that modulate HBV infection, including innate immune sensors and downstream interferon signaling components, and validate the HBV entry receptor NTCP *in vivo* in humanized mice. As a complementary approach, we also developed lentiviral vector-mediated expression of Cas9 and sgRNA, which provides sustained editing capacity suitable for experiments requiring stable perturbation or selection-based enrichment, albeit with an increased risk of cumulative off-target effects due to genomic integration. This framework bridges *in vitro* perturbation with *in vivo* validation in physiologically relevant human hepatocytes, enabling systematic study of liver biology and disease mechanisms.

## Results

### Lipofection-based RNP delivery to mpPHH

Existing RNP transfection protocols for cell lines are not directly applicable to primary cells. We therefore developed conditions tailored for primary human hepatocytes, using a non-targeting (NT) sgRNA as a negative control to assess baseline transfection toxicity and a commercially available lethal sgRNA pool (**Supplementary Table 1**) as a positive control for successful RNP delivery. Cell viability was quantified three days post-transfection using a fluorescence-based assay (PrestoBlue), allowing us to simultaneously assess transfection-induced toxicity (NT wells) and delivery efficiency (lethal sgRNA wells). We transfected mpPHH under various conditions by titrating the DharmaFECT Duo (Horizon Discovery) transfection reagent, the recombinant Cas9-to-sgRNA ratio, and the RNP concentration (**Supplementary Fig. 1a**). While direct quantification of transfection efficiency in mpPHH is challenging, we optimized conditions to maximize functional delivery while minimizing cytotoxicity, using the lethal sgRNA control as a surrogate readout for editing efficiency. We consistently observed near-complete loss of viability under optimized conditions, indicating highly efficient RNP uptake.

Based on reports that centrifugation enhances liposome-mediated transfection in other models^23^, we investigated whether centrifuging mpPHH immediately after applying the transfection mix containing RNP and DharmaFECT Duo would improve delivery. Centrifugation at 1,000 × g for one hour reduced cell viability in mpPHH transfected with lethal sgRNAs three days post-transfection (**Supplementary Fig. 1b**), indicating successful RNP delivery and functional gene editing. Notably, without centrifugation, lethal sgRNAs did not affect cell viability, suggesting minimal RNP uptake under these transfection conditions. Therefore, RNP delivery by lipofection in mpPHH critically depends on the centrifugation step. To optimize this step, we tested different centrifugation conditions, but further increases in either duration or g-force did not improve delivery efficiency (**Supplementary Fig. 1c**). These results establish an effective RNP lipofection protocol for mpPHH that includes centrifugation at 1,000 × g for one hour (**Fig. 1a**). To evaluate the reproducibility and generalizability of the platform, we compared RNP-mediated editing in two independently generated mpPHH cultures (from two humanized mice, #3760 and #3773) derived from the same PHH donor and in unpassaged PHH obtained from a different donor. Although successful editing was achieved in both PHH and mpPHH, as shown by the effects of transfecting lethal sgRNAs and *CYP3A4*-targeting sgRNAs, mpPHH consistently demonstrated improved long-term viability and maintenance of hepatocyte morphology following transfection, particularly at later time points after RNP delivery (**Supplementary Fig. 2).** This difference is likely attributable to the limited lifespan and progressive functional decline previously reported for conventional PHH cultures during prolonged *in vitro* maintenance^24^. Importantly, editing efficiency was quantified in CYP3A4-targeted cells using Sanger sequencing followed by Inference of CRISPR Edits (ICE) analysis (Synthego), which demonstrated approximately 96% editing efficiency in CYP3A4 sgRNA-transfected cells relative to NT controls. These findings are consistent with our optimization strategy to maximize functional RNP delivery and achieve near-complete gene disruption in mpPHH cultures.

**Fig. 1:**
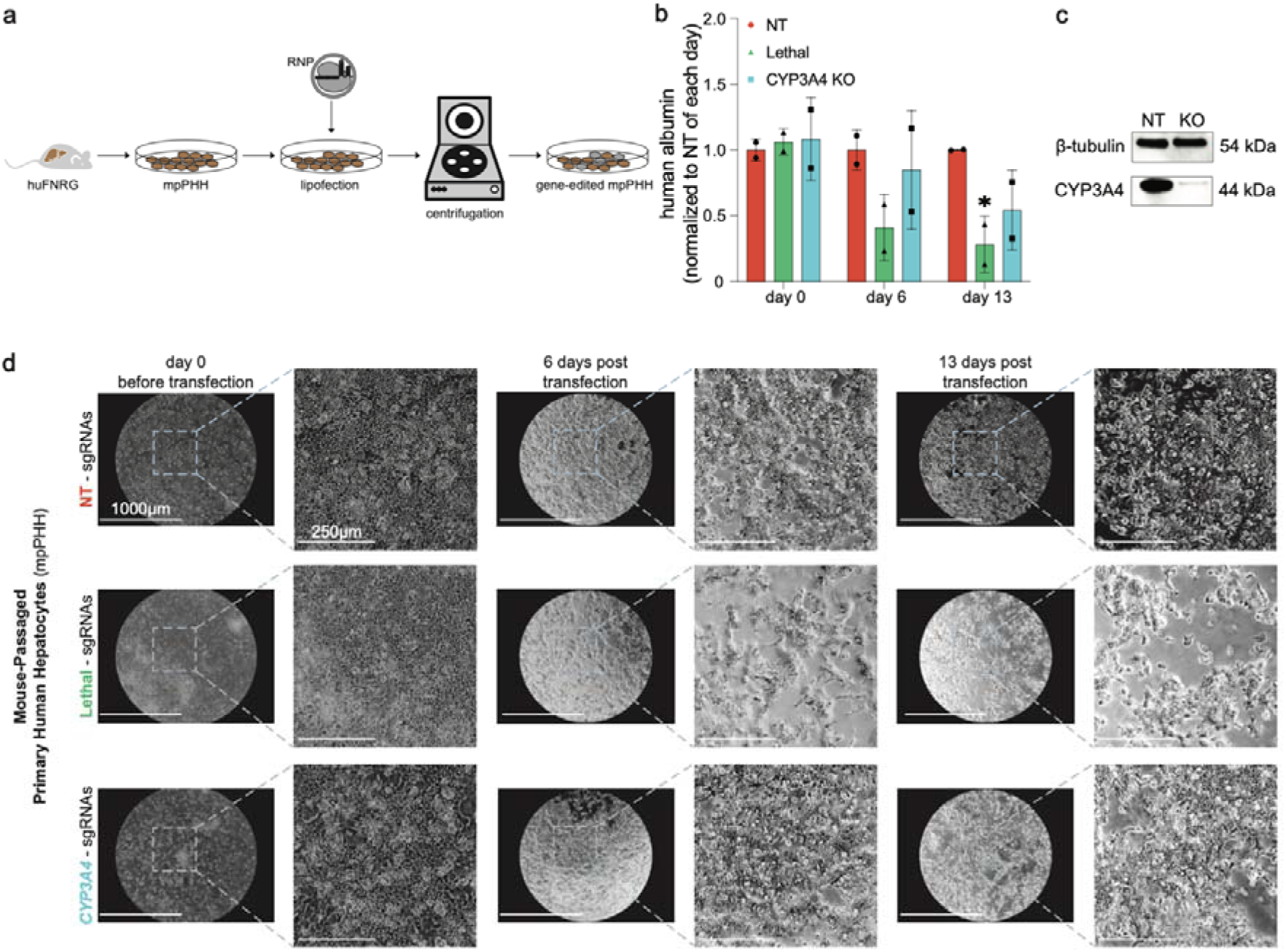
Lipofection-based RNP delivery supports long-term culture and gene editing in mpPHH. **a.** Schematic of the lipofection-based Cas9-sgRNA ribonucleoprotein (RNP) delivery workflow in mouse-passaged primary human hepatocytes (mpPHH) isolated from humanized huFNRG mice. mpPHH were transfected with RNP complexes by lipofection, followed by centrifugation-assisted delivery to generate gene-edited mpPHH cultures. **b.** Longitudinal human albumin secretion from mpPHH transfected with non-targeting (NT), lethal, or *CYP3A4*-targeting sgRNAs. Albumin levels were measured in culture supernatants and normalized to the NT condition at each corresponding time point. Data are shown as mean ± SD. **c.** Representative western blot showing CYP3A4 protein depletion in CYP3A4 KO vs NT mpPHH. β-tubulin served as the loading control. **d.** Representative bright-field microscopy images of mpPHH transfected with NT, lethal, or *CYP3A4*-targeting sgRNAs at day 0, day 6, and day 13 post-transfection. Overview images and magnified regions are shown for each condition and time point. NT and CYP3A4 KO mpPHH cultures maintained hepatocyte morphology over time, whereas lethal sgRNA-transfected cultures showed progressive loss of viability. Scale bars, 1000 μm for overview images and 250 μm for magnified images.

### Optimized transfection maintains mpPHH viability and function

Lipofection can cause significant cell toxicity^25^, which in non-dividing PHH may severely limit experimental applicability. To evaluate the impact of our optimized RNP delivery protocol on hepatocyte health, we monitored human albumin (hAlb) secretion, an established marker of hepatocyte function and differentiation^26,27^, in mpPHH transfected with NT, *CYP3A4*-targeting, or lethal sgRNAs (**Supplementary Table 1**) (**Fig. 1b**). hAlb levels were measured longitudinally throughout the experiment and normalized to NT controls at each time point. NT and CYP3A4 knockout (KO) mpPHH maintained hAlb secretion near baseline levels throughout the culture period, indicating that RNP delivery and CYP3A4 disruption do not substantially impair hepatocyte viability or function. In contrast, lethal sgRNA-transfected mpPHH showed a progressive decline in hAlb secretion, consistent with efficient delivery of RNP complexes and disruption of essential cellular genes. To assess gene-editing efficiency, mpPHH and unpassaged PHH were transfected with either NT sgRNAs or *CYP3A4*-targeting sgRNAs. Sanger sequencing followed by ICE analysis confirmed efficient editing of the *CYP3A4* locus, with 96% efficiency in *CYP3A4*-targeted mpPHH, whereas NT controls showed no detectable editing. Consistent with these DNA-level results, western blot analysis performed 15 days post-transfection demonstrated a marked reduction in CYP3A4 protein expression in edited mpPHH (**Fig. 1c**). Bright-field microscopy images were acquired at each medium change throughout the experiment, and hAlb levels were measured longitudinally at every corresponding time point (**Supplementary Fig. 2**; absolute values shown in **Supplementary Fig. 3**). Representative images shown in **Fig. 1d** correspond to the day of transfection (day 0), an intermediate time point (day 6 post-transfection), and the endpoint of the experiment (day 13 post-transfection), while the complete image series and all longitudinal hAlb measurements are provided in **Supplementary Fig. 2**.

To characterize the extended impact of our transfection protocol on hepatocyte function and morphology, we performed longitudinal hAlb measurements and imaging every 2-3 days across a two-week time course following transfection with NT or lethal sgRNAs (**Supplementary Fig. 2**). NT sgRNA-transfected mpPHH maintained hAlb secretion at near-baseline levels throughout the two-week time course, with only a modest decrease at endpoint, and retained their characteristic hepatocytic phenotype of polygonal cell shape and binucleated appearance^28^. In contrast, lethal sgRNA-transfected mpPHH showed a progressive loss of hAlb secretion, falling approximately fourfold below the pre-transfection baseline by two weeks post-transfection, with morphological deterioration first apparent by day 6 and progressing to a substantial loss of viable cells by the two-week endpoint. These longitudinal patterns were reproduced in mpPHH derived from two independent humanized mice, demonstrating that the protocol maintains cell function and morphology across biological replicates for at least two weeks in culture.

### Optimized lipofection protocol is applicable to unpassaged PHH

Although mpPHH offer substantial advantages in availability and scalability, unpassaged PHH remain widely used across the hepatocyte research community. To determine whether our optimized lipofection protocol generalizes beyond mpPHH, we applied identical transfection conditions, including DharmaFECT Duo concentration, Cas9-to-sgRNA ratio, and centrifugation at 1,000 × g for one hour, to unpassaged PHH and performed parallel longitudinal characterization (**Supplementary Fig. 2a**). Lethal sgRNA-transfected unpassaged PHH showed substantial loss of cell viability by day 6 post-transfection, whereas NT sgRNA-transfected cells maintained hAlb secretion and hepatocyte morphology at near-baseline levels throughout the two-week time course. These results establish that our protocol is compatible with both mpPHH and unpassaged PHH, broadening its applicability across common PHH culture systems. Notably, when compared on an absolute (non-normalized) basis, unpassaged PHH reached peak hAlb secretion comparable to mpPHH by day 4 post-transfection, but this secretion was not sustained: hAlb levels in unpassaged PHH declined toward baseline by the day 13 endpoint across all conditions, including NT controls, whereas mpPHH continued to secrete hAlb at substantially higher absolute levels through the same time point (**Supplementary Fig. 3**). This difference in the durability of hepatocyte function over the culture period is consistent with mpPHH’s greater suitability for screening applications that require a sustained functional window.

### CRISPR-Cas9 gene editing of *CYP3A4* disrupts protein expression and function

Having established efficient RNP delivery, we investigated whether lipofection-based CRISPR gene editing produces biologically relevant phenotypes in mpPHH, using the hepatocyte-relevant target *CYP3A4*. CYP3A4 is the most abundant cytochrome P450 enzyme and plays a central role in hepatic xenobiotic metabolism, converting hydrophobic compounds into more polar, water-soluble metabolites^29,30^. This phase I biotransformation^31^ metabolizes more than 30% of clinically used therapeutics^32^. CYP3A4 activity varies widely across the human population, complicating preclinical drug evaluations. Although PHH from different donors capture this diversity, the resulting variability complicates controlled comparisons between experimental conditions. Therefore, permanently disrupting *CYP3A4* in PHH from a single donor provides a controlled system for studying drug metabolism and pharmacokinetics^33^.

Before disrupting *CYP3A4*, we confirmed that our mpPHH system preserves CYP3A4 enzymatic function. Since dimethyl sulfoxide (DMSO) helps primary hepatocytes maintain differentiation during extended culture^34,35^, we assessed its effect on CYP3A4 phase I enzymatic activity using a luciferase-based assay (**Supplementary Fig. 4a**). Enzymatic activity declined rapidly without DMSO, whereas 2% DMSO maintained stable activity for at least 15 days. Immunofluorescence staining corroborated these findings, showing higher CYP3A4 expression in DMSO-treated mpPHH (**Supplementary Fig. 4b**). We therefore included 2% DMSO in all subsequent experiments.

To determine whether reduced protein expression translated into loss of enzymatic function, we performed a longitudinal analysis of CYP3A4 activity in mpPHH. Consistent with the reduction in protein levels (**Fig. 2a,b**), CYP3A4 enzymatic activity declined rapidly following gene editing and was significantly reduced by day 5 post-transfection (**Fig. 2c**). The delayed reduction in CYP3A4 activity is consistent with the largely non-proliferative nature of primary hepatocytes, where pre-existing CYP3A4 protein must be degraded before the full functional consequences of gene disruption become apparent. As a result, the effects of CYP3A4 knockout become increasingly evident several days after editing as existing protein pools undergo turnover. CYP3A4 activity continued to decrease throughout the experiment, reaching more than 75% reduction relative to NT controls by day 11, demonstrating efficient functional disruption of CYP3A4 in mpPHH.

**Fig. 2:**
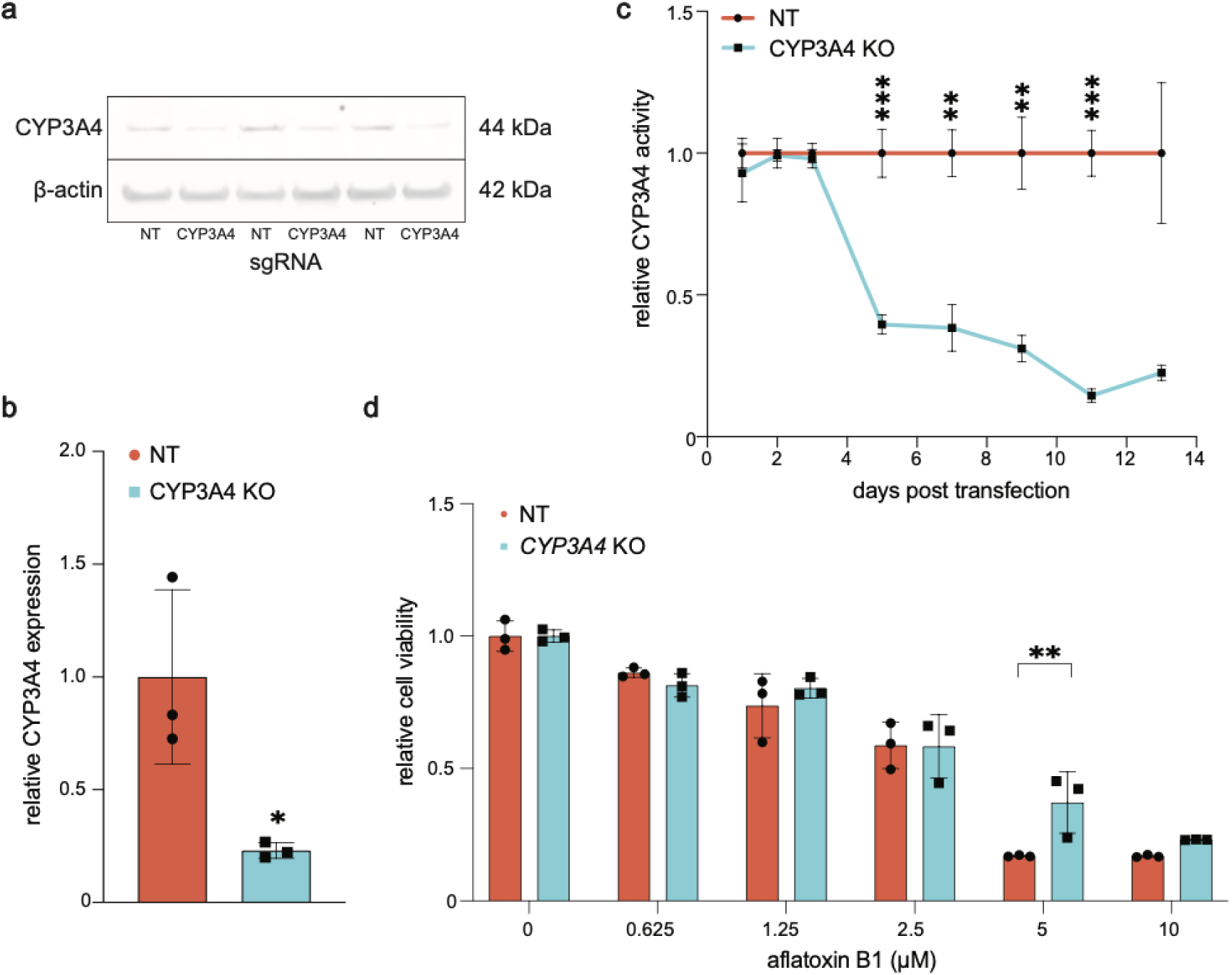
Lipofection-based RNP-mediated *CYP3A4* disruption demonstrates functional gene editing in mpPHH. **a.** Representative western blot analysis of CYP3A4 protein expression in NT vs CYP3A4 KO mpPHH. Three independent biological replicates are shown per condition. β-actin served as the loading control. **b.** Densitometry analysis of CYP3A4 protein expression from western blot in panel **a**. CYP3A4 levels were normalized to β-actin and are shown relative to NT controls. Data are mean ± SD from three biological replicates. **c.** Longitudinal CYP3A4 enzymatic activity following RNP transfection in mpPHH transfected with NT or CYP3A4-targeting sgRNAs. Activity was measured at days 1, 3, 5, 7, 9, 11, and 13 post-transfection using the luciferin-IPA substrate and is shown normalized to the time-matched NT sgRNA control to control for any baseline drift in CYP3A4 activity across the culture period; absolute CYP3A4 activity in non-edited mpPHH under identical DMSO-supplemented conditions is shown in Supplementary Fig. 4a. **d.** Functional assessment of *CYP3A4* disruption using aflatoxin B1 sensitivity. NT and CYP3A4 KO mpPHH were exposed to increasing concentrations of aflatoxin B1, and relative cell viability was measured. CYP3A4 KO cells showed reduced sensitivity to aflatoxin B1-induced toxicity, consistent with impaired CYP3A4-dependent bioactivation. Data are shown as mean ± SD. Statistical significance was assessed as indicated in Methods. *P < 0.05, **P < 0.01, ***P < 0.001.

To test whether *CYP3A4* disruption affects the metabolism of a clinically relevant xenobiotic, we challenged *CYP3A4*-edited mpPHH with aflatoxin B1 (AFB1). AFB1 is a mycotoxin produced by *Aspergillus* fungi that causes both acute and chronic liver injury and is a significant risk factor for hepatocellular carcinoma (HCC). AFB1 requires CYP3A4-mediated bioactivation to exert hepatotoxicity^36^, and silencing *CYP3A4* has been shown to reduce AFB1 cytotoxicity in PHH^37^. We hypothesized that Cas9-mediated disruption of *CYP3A4* would partially protect mpPHH from AFB1’s cytotoxic effects. Seven days after transfection with NT or *CYP3A4*-targeting RNPs, we exposed mpPHH to different concentrations of AFB1. AFB1 decreased cell viability in a dose-dependent manner in NT sgRNA-transfected mpPHH (**Fig. 2d**). *CYP3A4* KO preserved mpPHH viability after AFB1 treatment, demonstrating partial protection from AFB1-induced cytotoxicity. The residual toxicity seen at 10 µM AFB1 is consistent with AFB1 bioactivation through *CYP3A4*-independent pathways and with residual CYP3A4 protein in incompletely edited cells.

These findings establish that lipofection-based CRISPR-Cas9 gene editing in mpPHH produces functionally relevant phenotypes and may be leveraged to study xenobiotic metabolism in a controlled hepatocyte model.

### *CYP3A4*-edited mpPHH engraft and expand in human liver chimeric mice

Although mpPHH cannot proliferate in cell culture, they can be expanded in FNRG mice^12^. To determine whether gene-edited mpPHH retain this capacity, we transplanted NT or *CYP3A4*-targeted mpPHH into FNRG mice, as described previously^12,38^ (**Fig. 3a**). We tracked liver humanization by measuring serum hAlb levels over time. Both NT and *CYP3A4* KO groups achieved high, stable hAlb levels (**Fig. 3b**), demonstrating that our editing protocol did not impair engraftment or subsequent expansion of mpPHH in FNRG mice.

**Fig. 3:**
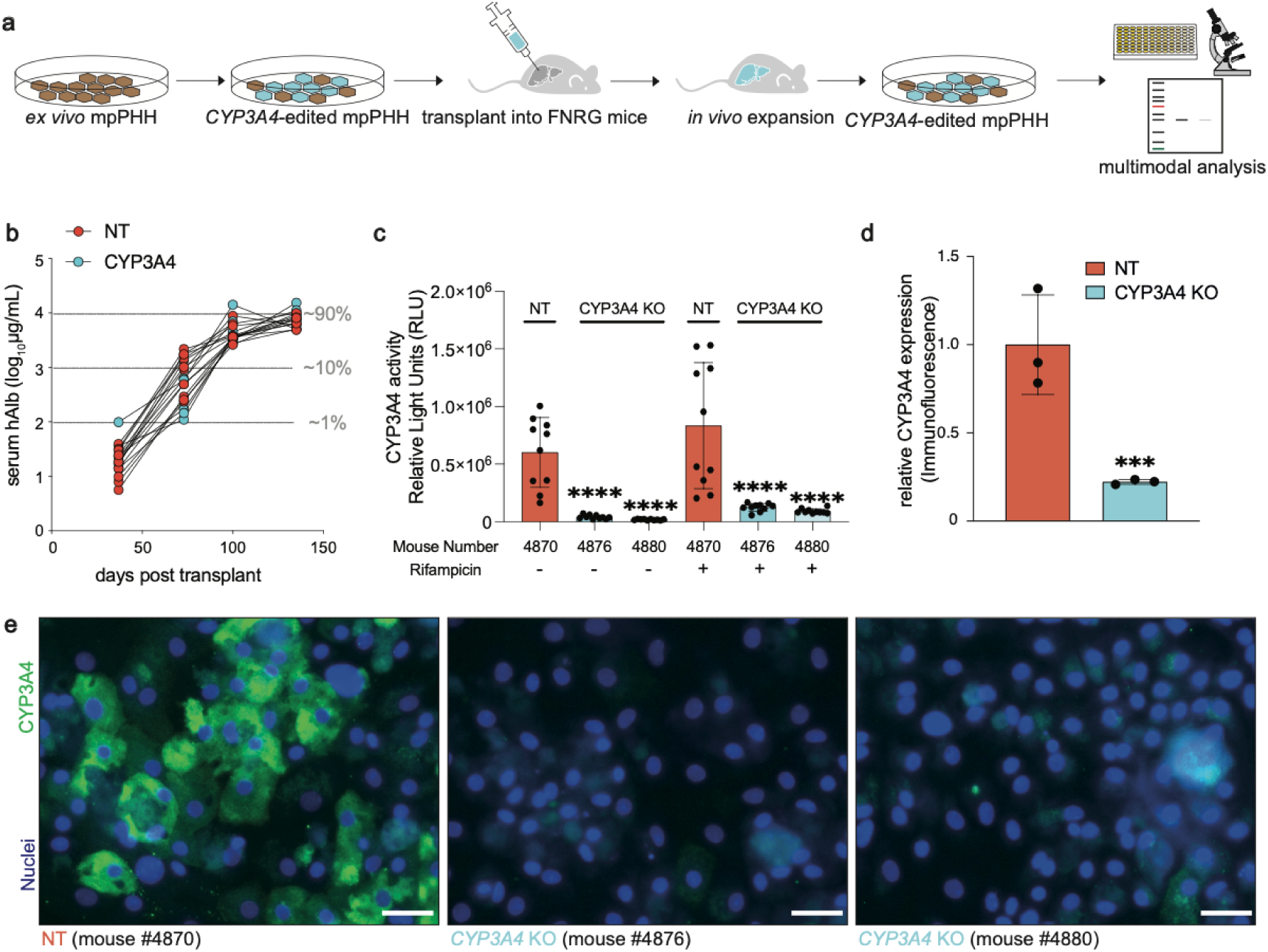
CYP3A4 KO mpPHH engraft in FNRG mice and retain the edited phenotype after *in vivo* expansion. **a.** Schematic of the workflow used to transplant CYP3A4 KO mpPHH into FNRG mice, expand edited human hepatocytes *in vivo*, re-isolate mpPHH, and perform multimodal analysis. **b.** Longitudinal serum human albumin levels in FNRG mice transplanted with NT or CYP3A4 KO mpPHH. Serum hAlb is shown on a log_10_ scale as a marker of human hepatocyte engraftment and expansion. Approximate thresholds corresponding to ∼1%, ∼10%, and ∼90% humanization are indicated. **c.** CYP3A4 enzymatic activity in re-isolated mpPHH from one NT-engrafted mouse and two CYP3A4 KO-engrafted mice. Cells were cultured with vehicle or rifampicin, and CYP3A4 activity was measured using the luciferin-IPA substrate. Rifampicin induced CYP3A4 activity in NT cells, whereas CYP3A4 KO-derived cells showed markedly reduced activity while retaining some responsiveness to induction. **d.** Quantification of CYP3A4 protein expression in re-isolated mpPHH by immunofluorescence. Data are shown as mean ± SD. **e.** Representative immunofluorescence images of re-isolated mpPHH stained for CYP3A4 and nuclei. Images are shown from one NT mouse and two CYP3A4 KO mice. Scale bars, 50 μm.

To confirm that the edited phenotype was maintained after in vivo expansion, we re-isolated mpPHH from humanized mice, including one mouse engrafted with NT control mpPHH and two mice engrafted with CYP3A4 KO mpPHH. After three days in culture, CYP3A4 enzymatic activity was significantly reduced in mpPHH derived from CYP3A4-edited grafts (**Fig. 3c**). Treatment with rifampicin increased CYP3A4 activity by approximately eight-fold in NT control cells. Although CYP3A4 KO cells exhibited substantially lower basal activity, they retained responsiveness to rifampicin stimulation. Immunofluorescence analysis further confirmed at least a four-fold reduction in CYP3A4 protein expression in edited cells relative to controls (**Fig. 3d,e**). Together, these findings demonstrate that CYP3A4 KO mpPHH can be expanded in chimeric mice while maintaining a stable edited phenotype.

### CRISPR-Cas9 gene editing in mpPHH enables arrayed screening for HBV host factors

Next, we tested whether our lipofection-based CRISPR protocol could be leveraged for high-throughput screening in mpPHH. We obtained a 96-well synthetic sgRNA library (**Supplementary Table 1**) containing 85 genes (4 sgRNAs per gene). NT and lethal sgRNAs were included as negative and positive controls, respectively. These 85 genes were selected for their relevance to innate immunity and liver biology, or for their prior identification as HBV-related host factors. MpPHH were seeded into three replicate 96-well plates and, three days later, transfected with RNP complexes, following our optimized protocol (**Fig. 4a**). Successful RNP delivery was confirmed by observing cell deterioration in lethal sgRNA-transfected wells. On day five post-transfection, mpPHH were infected with HBV. Seven days post-infection, we measured HBV surface antigen (HBsAg) levels in culture supernatants using a chemiluminescence immunoassay (CLIA).

**Fig. 4:**
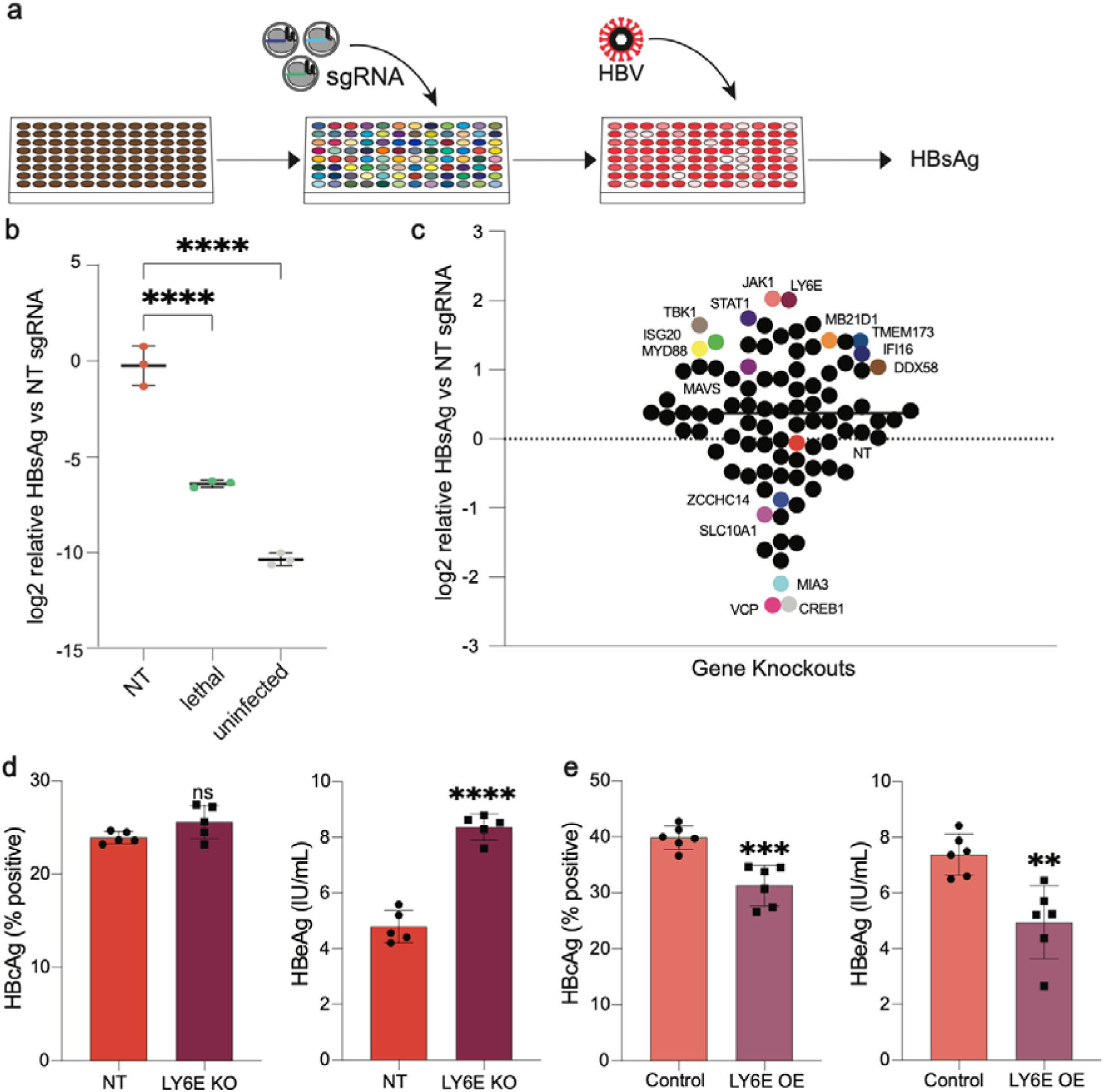
Arrayed CRISPR screening in mpPHH identifies host factors that modulate HBV infection. **a.** Schematic of the arrayed CRISPR screening workflow in mpPHH. Cells were transfected with RNP complexes in an arrayed format, infected with HBV, and culture supernatants were analyzed for HBsAg. **b.** Log_2_-transformed relative HBsAg levels in culture supernatants from HBV-infected mpPHH transfected with NT or lethal sgRNAs, and from uninfected NT control wells. HBsAg values were normalized to HBV-infected NT controls before log_2_ transformation. **c.** Arrayed CRISPR screen showing log_2_ relative HBsAg levels for each targeted gene knockout compared with NT controls. Each point represents a gene-targeting condition. Candidate hits are labeled, including genes whose disruption increased or decreased HBsAg levels relative to NT controls. **d.** Validation of LY6E knockout in HBV-infected HepG2-NTCP cells. HBcAg-positive cells were quantified by immunofluorescence, and HBeAg levels in culture supernatants were measured by CLIA. **e.** Validation of LY6E overexpression in HBV-infected HepG2-NTCP cells. HBcAg-positive cells were quantified by immunofluorescence, and HBeAg levels were measured by CLIA. Data are shown as mean ± SD. Statistical significance was assessed as indicated in Methods. ns, not significant; **P < 0.01, ***P < 0.001, ****P < 0.0001.

Evaluating negative and positive controls revealed robust assay performance (**Fig. 4b**). HBsAg levels differed by three orders of magnitude between NT sgRNA-transfected HBV-infected cells and uninfected controls. Lethal sgRNA-transfected cells had HBsAg two orders of magnitude lower than NT. Together, these confirmed successful RNP delivery followed by efficient HBV infection (**Fig. 4b**). Further, these results demonstrate adequate signal-to-noise ratios for screening applications. This highlights that mpPHH can be efficiently transfected and edited, and can also be further manipulated downstream.

Our screens identified multiple genes affecting HBV infection (**Fig. 4c**). As expected, *VCP* disruption, included as a cell death control consistent with the essential pro-survival role of VCP^39^, produced the lowest HBsAg levels. Known HBV host factors showed decreased infection upon disruption, including *ZCCHC14* (essential for HBsAg production^40^) and *SLC10A1* (encoding the entry receptor NTCP^41^). Notably, disruption of previously uncharacterized genes also reduced HBV infection, such as *MIA3*, whose protein product, TANGO1, mediates COPII-dependent ER-to-Golgi trafficking^42^.

Strikingly, disruption of multiple innate immune pathway components enhanced HBV infection. Genes in the cGAS-STING pathway showed increased HBV levels when disrupted, consistent with knockdown data in hepatoma cell lines^43^. Similarly, disrupting the RIG-I pathway (*DDX58/RIG-I*, *MAVS*, *TBK1*), which senses HBV pregenomic RNA and triggers interferon signaling^44^, enhanced infection. Downstream interferon signaling components (*JAK1*, *STAT1*) and interferon-stimulated genes with known anti-HBV activity (*MYD88*^45^, *ISG20*^46^) also showed increased HBV levels upon disruption, confirming successful perturbation of antiviral pathways.

Statistical analysis using nested one-way ANOVA with Benjamini-Krieger-Yekutieli correction for multiple comparisons (FDR = 0.05) identified only *JAK1* and *LY6E* as statistically significant hits (adjusted p = 0.015 and p = 0.017, respectively)^47^. We observed variability across screens, suggesting that future screens will benefit from increased replicate numbers to improve statistical power and precision.

To validate LY6E as a candidate antiviral host factor identified in the arrayed screen, we performed gain- and loss-of-function experiments in HepG2-NTCP cells. Overexpression of LY6E resulted in a significant reduction in HBV infection, as evidenced by decreased hepatitis B e antigen (HBeAg) secretion and reduced intracellular hepatitis B core antigen (HBcAg) levels compared to control cells (**Fig. 4e**). Conversely, CRISPR-Cas9-mediated knockout of *LY6E* led to a marked increase in both HBeAg and HBcAg levels, consistent with enhanced viral replication (**Fig. 4d**). These reciprocal phenotypes support a restrictive role for LY6E in HBV infection. Together, these data independently corroborate the screening results and establish LY6E as a functional antiviral factor in hepatocyte-derived cell lines.

Collectively, these data show that CRISPR-Cas9 editing in mpPHH supports arrayed screening approaches and can identify both known and putative HBV host factors, providing a platform for systematic study of hepatocyte-pathogen interactions.

### *SLC10A1*-edited mpPHH protect chimeric mice from HBV infection and validate screening hits *in vivo*

Having validated our approach with *CYP3A4* and confirmed *SLC10A1* as an expected positive control in our arrayed screen, we sought to demonstrate that our platform enables seamless progression from *in vitro* screening to *in vivo* validation. *SLC10A1* encodes NTCP, the functional receptor for HBV entry, making it an attractive target for proof-of-concept validation (**Fig. 5a**). We engrafted mpPHH transfected with RNP complexes targeting *SLC10A1* or NT control sgRNAs into FNRG mice and monitored serum hAlb levels to assess the degree of liver humanization. Once robust humanization was established (**Fig. 5b**), typically 8-12 weeks post-transplantation, mice were challenged with HBV. Serum HBV DNA levels were subsequently monitored longitudinally by qPCR for 130 days (**Fig. 5c**). Three NTCP-KO mice (#3404, #3411, #4350) either died spontaneously or were euthanized per humane endpoint criteria (severe lethargy and/or weight loss exceeding 25% of body weight unresponsive to NTBC rescue) and were therefore not followed through the full 130-day observation period used for the remaining animals. In the control groups, two uninfected mice (#4059, #4066) were last measured at day 86, and three HBV-infected control mice (#4055, #4067, #4069) were euthanized for planned liver harvest after reaching peak viremia to generate HBV-infected mpPHH. All animals that exited the study before day 130, and their respective time points, are indicated in **Supplementary Fig. 5**. The study included three groups: mice engrafted with *SLC10A1*-edited mpPHH (NTCP KO huFNRG, n = 6), HBV-challenged control humanized mice (huFNRG, n = 6), and unchallenged humanized controls (n = 3). Consistent with the essential role of NTCP in HBV entry, mice engrafted with *SLC10A1*-edited mpPHH exhibited substantially reduced viral loads compared with HBV-challenged controls. While some variability in HBV titers was observed within the control group, this is expected in humanized mouse models and likely reflects differences in engraftment efficiency and donor-derived hepatocyte functionality. Importantly, humanization levels, as assessed by serum hAlb, were comparable between groups, indicating that differences in viral load are not driven by unequal engraftment.

**Fig. 5:**
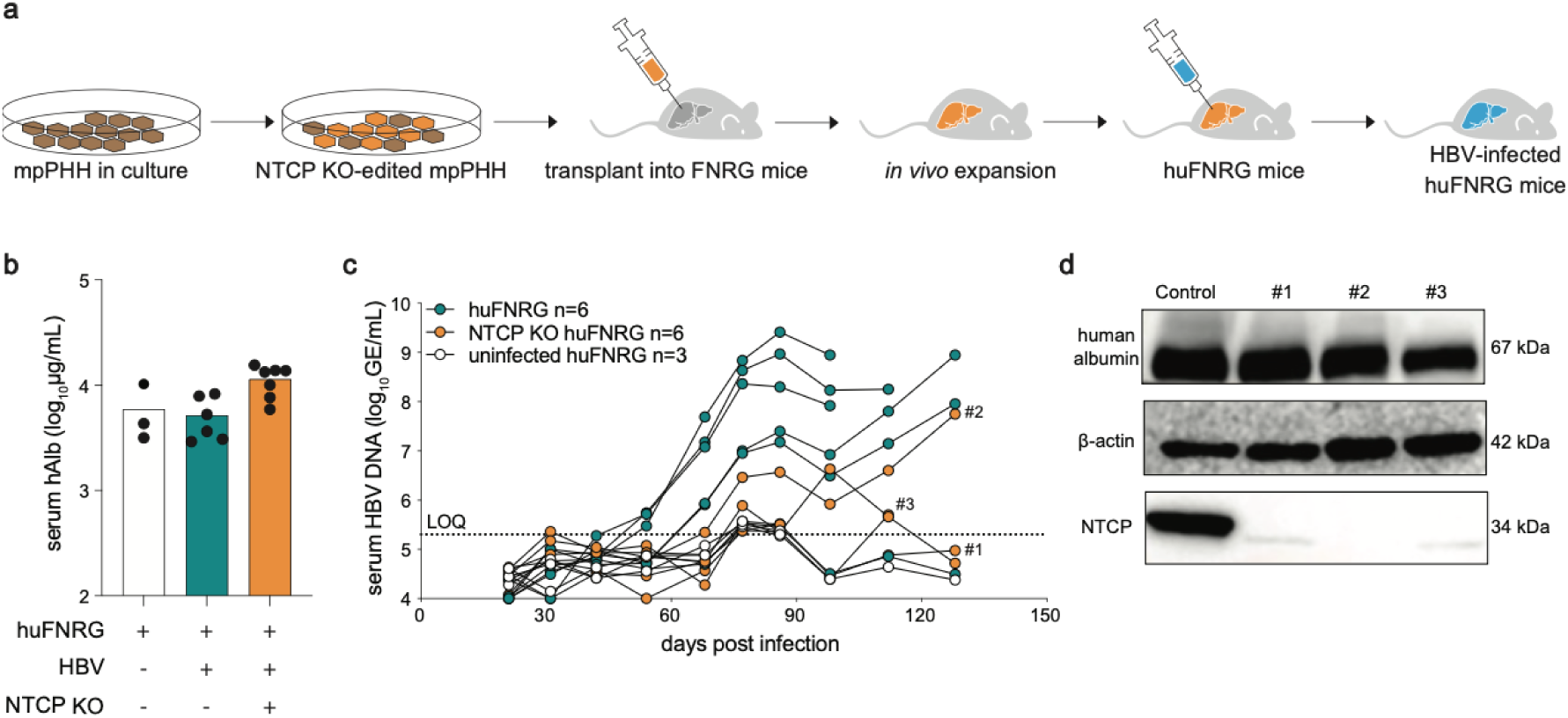
NTCP KO mpPHH reduce HBV infection after transplantation into humanized FNRG mice. **a.** Schematic of the workflow for *SLC10A1*/NTCP editing in mpPHH followed by transplantation into FNRG mice, *in vivo* expansion, humanization, and HBV challenge. **b.** Serum human albumin levels in humanized FNRG mice before HBV challenge. Groups include uninfected huFNRG mice, HBV-infected huFNRG mice, and HBV-infected mice engrafted with NTCP KO mpPHH. Serum human albumin (hAlb) is shown on a log_10_ scale. **c.** Longitudinal serum HBV DNA levels following HBV challenge. HBV DNA was quantified by qPCR over time in control huFNRG mice, NTCP KO huFNRG mice, and uninfected huFNRG controls. The dotted line indicates the assay limit of quantification. Mice that did not reach the full 130-day study endpoint either died spontaneously, were euthanized per IACUC-approved humane criteria, or were euthanized for planned liver harvest after reaching peak viremia; individual animal trajectories, including the time point at which each of these mice exited the study and the reason for each exit, are shown in **Supplementary** Fig. 5. **d.** Western blot analysis of NTCP expression in hepatocytes re-isolated from humanized mice at the experimental endpoint. Human albumin served as a marker of human hepatocytes, and β-actin served as a loading control. Each lane represents cells re-isolated from an individual mouse.

Human hepatocyte engraftment kinetics in our study were consistent with previous reports^12,48^ using FNRG mice, with stable humanization achieved within 8-12 weeks post-transplantation. This suggests that CRISPR-mediated disruption of *SLC10A1* does not impair hepatocyte engraftment or expansion *in vivo*. Collectively, these data demonstrate that our platform supports robust in vivo validation of screening hits in a physiologically relevant model.

After 130 days, we re-isolated hepatocytes from chimeric mice to verify editing efficiency. Western blot analysis confirmed robust knockdown of the NTCP protein in *SLC10A1*-edited samples, with hAlb serving as a human hepatocyte marker and loading control. High hAlb signal in all samples was consistent with sustained high levels of liver humanization at the experimental endpoint (**Fig. 5d**). These results establish a complete workflow from *in vitro* CRISPR screening to *in vivo* validation in humanized mice, highlighting the robustness of our platform.

### Lentiviral vector-mediated CRISPR-Cas9 delivery provides a complementary approach for gene disruption in mpPHH

While lipofection of recombinant Cas9-sgRNA RNP complexes provides an effective method for gene perturbation in mpPHH, we also evaluated a complementary strategy based on lentiviral vector-mediated expression of Cas9 and sgRNAs. In this approach, mpPHH were co-transduced with two lentiviral vectors (**Fig. 6a**). Specifically, we co-transduced mpPHH with a lentivirus carrying four sgRNAs^49^ targeting *SLC10A1* (NTCP) (**Supplementary Table 1**) together with a separate lentiviral Cas9-RFP vector that we engineered in a pSCRPSY backbone (GenBank: KT368137.1). After 14 days of permitting protein turnover, western blot analysis confirmed a substantial reduction in NTCP expression (**Fig. 6b**). Unlike the lipofection experiments, in which pre-assembled Cas9-sgRNA RNPs were delivered directly to cells, the lentiviral approach relies on intracellular expression of Cas9 and sgRNAs after transduction. A parallel approach targeting *CYP3A4* using lentiviruses encoding sgRNAs^49^ (**Supplementary Table 1**) and Cas9 under blasticidin selection (Cas9-Blast) was also employed (**Fig. 6c**). Cas9 was delivered using lentiviral vectors expressing Cas9 under constitutive promoters. Specifically, the Cas9-RFP construct, driven by a CMV promoter, was used to enable visual assessment of transduction efficiency, whereas a separate Cas9-blasticidin construct, driven by an EF1α promoter, was used for selection-based enrichment of Cas9-expressing cells. The RFP-expressing system allowed direct visualization of transduced cells and rapid qualitative assessment of delivery efficiency. In parallel, the blasticidin-selectable Cas9 construct enabled enrichment of edited cell populations, which was particularly important for experiments requiring sustained Cas9 expression and for downstream applications such as transplantation into mice, where a higher proportion of edited cells is desirable. We observed approximately an 80-85% reduction in NTCP levels and about 35-40% in CYP3A4, as measured by western blot. These differences likely reflect gene- and sgRNA-dependent variation in editing efficiency and/or protein turnover.

**Fig. 6:**
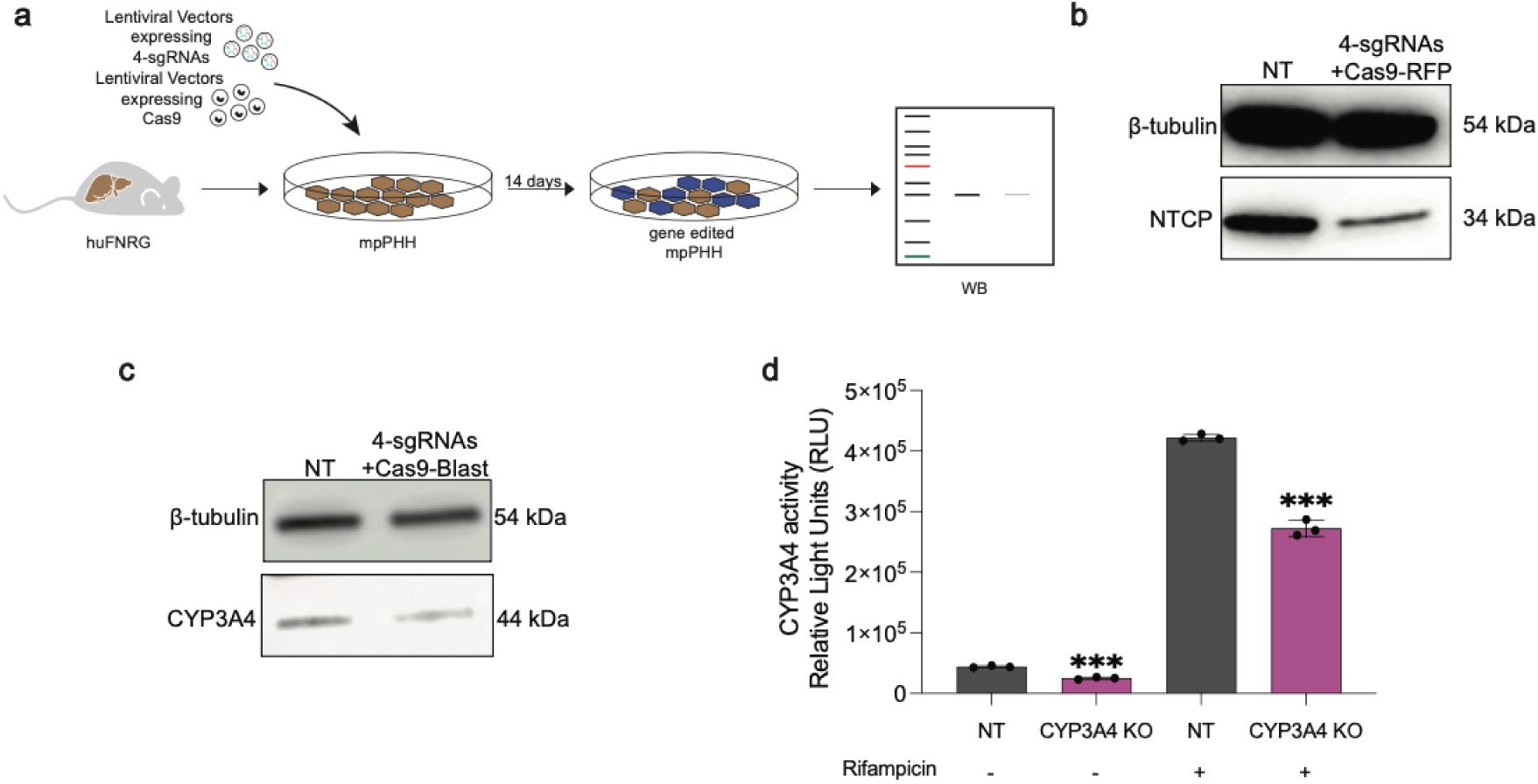
Lentiviral CRISPR-Cas9 delivery enables gene disruption in mpPHH. **a.** Schematic of the lentiviral CRISPR-Cas9 workflow in mpPHH. Cells were co-transduced with lentiviral vectors expressing Cas9 and lentiviral vectors expressing four sgRNAs targeting the gene of interest, followed by analysis 14 days post-transduction. **b.** Western blot analysis of NTCP protein expression in mpPHH transduced with NT control vectors or *SLC10A1*-targeting sgRNAs together with Cas9-RFP. β-tubulin served as the loading control. **c.** Western blot analysis of CYP3A4 protein expression in mpPHH transduced with NT control vectors or *CYP3A4*-targeting sgRNAs together with Cas9-Blast. β-tubulin served as the loading control. **d.** CYP3A4 enzymatic activity in NT and CYP3A4 KO mpPHH cultured with or without rifampicin. Activity was measured using the luciferin-IPA substrate and is shown as relative light units. Data are shown as mean ± SD. Statistical significance was assessed as indicated in Methods. ***P < 0.001.

To evaluate the functional effects of *CYP3A4* disruption, we treated mpPHH with rifampicin, a well-known inducer of CYP3A4 expression and activity. Although *CYP3A4* KO cells showed lower baseline activity, they still responded to rifampicin induction (**Fig. 6d**). This residual inducibility likely reflects incomplete *CYP3A4* disruption in the lentiviral setting, potentially due to heterogeneous transduction efficiency and/or variable sgRNA-mediated editing efficiency.

Together, these data show that lentiviral vector-mediated delivery of Cas9 and sgRNAs can achieve functionally relevant gene disruption in mpPHH and provide a complementary alternative to direct RNP lipofection. Whereas RNP lipofection is rapid and well-suited for short-term perturbation and arrayed screening, lentiviral delivery may be useful in settings where prolonged expression or enrichment of edited cells is advantageous.

However, in contrast to transient RNP delivery, lentiviral approaches rely on genomic integration and sustained expression of Cas9 and sgRNAs, which may increase the risk of cumulative off-target editing events. Despite this, we did not observe cytotoxicity or loss of hepatocyte function under the conditions tested, suggesting that lentiviral delivery is well tolerated in mpPHH and can be applied for longer-term studies when stable gene perturbation is required^12,48,50^. Additionally, lentiviral delivery provides a scalable and cost-effective alternative for large-scale applications, as Cas9- and sgRNA-expressing viral stocks can be generated in bulk and used across experiments, reducing reliance on repeated preparation and transfection of recombinant Cas9 protein and synthetic sgRNAs required for RNP-based approaches.

## Discussion

In this study, we establish a scalable platform for CRISPR-Cas9-mediated gene disruption and functional interrogation in primary human hepatocytes. By leveraging mouse-passaged PHH (mpPHH), which can be expanded in liver-humanized mice, our approach overcomes key limitations of traditional PHH systems, including limited cell availability and restricted experimental scalability. Importantly, this system enables not only efficient *in vitro* gene editing but also the expansion and phenotypic validation of edited hepatocytes *in vivo*. Our data demonstrate that gene disruption in mpPHH results in sustained, functionally meaningful phenotypes, as shown by *CYP3A4* editing, which affects xenobiotic metabolism, and by *SLC10A1* targeting, which reduces HBV infection in humanized mice. In addition, we show that this platform supports arrayed CRISPR screening to identify host factors involved in hepatocyte biology and viral infection. Together, these findings position mpPHH-based CRISPR approaches as a versatile system for functional genomics studies in human hepatocytes, bridging *in vitro* perturbation with *in vivo* validation.

Methods for straightforward gene editing in PHH that support flexible experimental design and unbiased large-scale investigations have been lacking. In this study, we present one of the first protocols for efficient CRISPR-Cas9 gene editing and screening in cultured mpPHH. We developed an optimized lipofection workflow for delivering RNP that achieves editing efficiencies exceeding 90% while maintaining cell viability. The addition of a centrifugation step proved crucial for effective RNP uptake, a finding with broader implications for transfecting other non-dividing primary cell types. Importantly, edited mpPHH retained their differentiated phenotype, as shown by stable albumin secretion and preserved hepatocyte morphology over long-term culture periods, addressing a major concern related to genetic manipulation in primary hepatocytes.

Our validation using *CYP3A4* demonstrates the functional importance of our approach for studying drug metabolism, a fundamental aspect of hepatocyte biology. The ability to disrupt *CYP3A4* and observe corresponding changes in xenobiotic metabolism, specifically protection from AFB1 cytotoxicity, confirms our platform’s suitability for pharmacological studies. Importantly, *CYP3A4* KO mpPHH retained their edited phenotype after e^x^pansion through FNRG mice, providing a scalable solution to the quantity limitations that have historically plagued primary hepatocyte research. This capability enables investigators to generate panels of edited hepatocytes from a single donor, effectively modeling genetic variation in drug metabolism while controlling for donor-specific variables, a major advance in personalized medicine.

The successful deployment of arrayed CRISPR screening in mpPHH establishes a new approach in hepatocyte functional genomics. While pooled screens have transformed our understanding of cancer and other dividing cells, their application to non-dividing PHH has been limited. Our 85-gene proof-of-concept study demonstrates that arrayed screens can overcome this limitation. We identified expected hits like *SLC10A1* and *ZCCHC14*, as well as new candidates such as *LY6E*, confirming our platform’s sensitivity. LY6E, like several interferon-stimulated genes, exhibits context-dependent effects on viral infection: it promotes HIV-1 entry and replication^51^ and enhances internalization of flaviviruses including West Nile, Zika, and dengue virus^52^, and enhances infection by influenza A and other RNA viruses at a late entry step, after endosomal escape, through an evolutionarily conserved mechanism^53^. Conversely, LY6E potently restricts entry of multiple coronaviruses^54^, including SARS-CoV, SARS-CoV-2, and MERS-CoV, by impairing spike-mediated membrane fusion. Together, these findings position LY6E as a context-dependent modulator of viral infection rather than a classical restriction factor. In our HBV-HepG2-NTCP model, the consistent loss- and gain-of-function effects (**Fig. 4d,e**) support a restrictive role for LY6E in HBV infection. Furthermore, the predictable responses of innate immune components, such as increased infection following disruption of cGAS-STING, RIG-I/MAVS, and JAK/STAT pathways, validate our screening approach. These findings underscore the importance of intact innate immunity in controlling HBV in primary hepatocytes.

The variability observed between screening replicates emphasizes important considerations for future studies. Primary hepatocytes, even when derived from the same donor and expanded through mice, likely maintain intrinsic heterogeneity that contributes to experimental variability. Future screens should include additional replicates and, where appropriate, leverage the stable expression provided by our lentiviral platform for longer durations. Furthermore, using multiple phenotypic readouts, such as combining HBsAg measurements with imaging- and molecular-based analyses, could provide more robust hit identification.

Our demonstration that screening hits can be validated *in vivo* using humanized mice establishes a complete discovery pipeline from initial screening to preclinical validation. The protection provided by *SLC10A1* disruption in humanized mice not only confirms NTCP as the essential HBV entry receptor in a physiologically relevant model but also shows the therapeutic potential of targeting host factors. This seamless progression from *in vitro* discovery to *in vivo* validation addresses an important bottleneck in translational hepatology research.

The development of a complementary lentiviral delivery system expands the versatility of our platform. While RNP lipofection offers rapid, efficient editing suitable for most applications, lentiviral delivery provides advantages for experiments requiring stable Cas9 expression, multiplexed targeting, or selection-based enrichment. In addition, lentiviral-mediated expression of sgRNAs and Cas9 can be more cost-effective than using recombinant Cas9 and synthetic sgRNAs. This dual-platform approach ensures that our system can accommodate diverse experimental designs and research questions.

Our findings have immediate implications for multiple areas of liver research. For drug development, the ability to systematically knock out drug-metabolizing enzymes and transporters will accelerate understanding of drug-drug interactions and individual variation in drug response. For infectious disease research, our platform enables systematic identification of host factors required for hepatitis D virus (HDV), malaria, and other hepatotropic pathogens. For metabolic disease, CRISPR screening in mpPHH could identify novel regulators of lipid metabolism, glucose homeostasis, and protein synthesis. Furthermore, the ability to expand edited hepatocytes in mice opens the possibility of validating phenotypes *in vivo*.

Several limitations merit consideration. First, while our editing efficiency can exceed 90%, complete knockout may not be possible for all genes, which could limit the detection of subtle phenotypes. Second, the current throughput of arrayed screening, while suitable for focused libraries, cannot match the genome-wide scale achievable with pooled approaches. Third, the need for specialized expertise in both hepatocyte culture and mouse husbandry may limit widespread adoption. Future efforts should aim to improve editing efficiency, potentially through optimized RNP formulations or alternative delivery methods, scale up arrayed screening via automation to a 384-well format, and develop simplified protocols accessible to non-specialist laboratories.

In conclusion, we have developed lipofection- and lentiviral-based protocols for CRISPR- Cas9 gene editing in primary human hepatocytes that efficiently disrupt target genes in both single-gene and large-scale experiment settings, and demonstrate the utility of this approach through *in vivo* validation in humanized mice. This advance enables future studies to more effectively leverage the potential of primary human hepatocytes, facilitating the discovery of therapeutic targets for liver diseases, particularly chronic hepatitis B, which affects millions worldwide. As CRISPR technologies continue to evolve, including base editing, prime editing, and CRISPR activation/inhibition, our platform offers a solid foundation for applying these cutting-edge tools in the most physiologically relevant human hepatocyte model available.

## Methods

### Mice

*Fah*^-/-^ NOD *Rag1*^-/-^ *Il2rg*^null^ (FNRG) were humanized with PHH from a pediatric human donor (cat#HUM4188, Lonza) and from mpPHH isolated from humanized mice as previously described^12^. The use of de-identified PHH does not qualify as human subjects research and is therefore exempt from institutional review board review. FNRG mice were maintained on an ad libitum chow diet with 0.12% amoxicillin and drinking water containing 16 mg/mL NTBC (cat#20-0028, Yecuris). All procedures involving animals were approved by the Institutional Animal Care and Use Committees at Weill Cornell Medicine under protocol 2015-0031. The mice were group-housed (5 per cage) under a 12-hour light/dark cycle at 21°C and 50% relative humidity to ensure optimal environmental conditions. Preconditioning of the animals was performed using retrorsine^12^. To generate humanized mice with modified grafts, mpPHH isolated from humanized FNRG mice were modified in culture (gene knockout), and then the cells were detached and transplanted into FNRG mice. Once these mice were humanized based on serum hAlb levels, the mice were either sacrificed for mpPHH isolation (CYP3A4 KO mice) or for HBV infection studies (NTCP KO mice).

### Mouse-passaged primary human hepatocytes (mpPHH)

The primary hepatocytes used in this study were derived from human donors and expanded in liver-humanized mice to generate mpPHH. Unless otherwise indicated, experiments were performed using mpPHH derived from a single donor (cat#HUM4188, Lonza) to minimize variability and enable controlled comparisons between conditions. Biological replicates represent independent wells or independent mouse-derived hepatocyte preparations from the same donor, whereas technical replicates correspond to repeated measurements within the same experimental unit. While the use of mpPHH reduces donor-to-donor variability compared to unpassaged PHH, intrinsic heterogeneity may still contribute to variability in gene editing efficiency and phenotypic readouts. While the present study primarily utilized mpPHH derived from a single donor to minimize variability and enable controlled comparisons between experimental conditions, previous work using this platform successfully expanded and characterized hepatocytes from 13 independent human donors^12^, supporting the broader applicability and generalizability of the mpPHH system. Mouse-passaged primary human hepatocytes (mpPHH) were isolated from highly humanized FNRG mice as previously described^12^ and humanization levels were based on serum hAlb levels. In brief, humanized FNRG mice were anesthetized with ketamine/xylazine, and a 24-gauge angiocath was inserted into the inferior vena cava. Then, the portal vein was cut, and the mouse liver was perfused sequentially with PBS^−/−^ supplemented with heparin, Hanks buffered saline solution (HBSS) supplemented with 5 mM ethylenediaminetetraacetic acid (EDTA) and 50 mM 2-[4-(2-hydroxyethyl)piperazin-1-yl] ethanesulfonic acid (Hepes), and finally, HBSS supplemented with 0.05% Type IV collagenase (Sigma Aldrich) and 1 U/mL DNase (Thermo Fisher). After digestion, the liver was disrupted over a 70-μm cell strainer, and the cell suspension (50 mL) was centrifuged at 50 x g for 5 min at 4°C using an Allegra X-14R Centrifuge (Beckman Coulter). The supernatant was gently aspirated, and the cells were washed once with PBS^−/−^. The cell pellet was resuspended in 10 mL PBS^−/−^ and gently mixed with an equal volume of Percoll working solution (cat#17–0891-01, GE Healthcare). Percoll working solution consisted of 5.4% 10x PBS^−/−^, 48.6% Percoll, and 46% William’s E medium (WEM) (cat#12551–032; ThermoFisher). The cell suspension was centrifuged at 100 x g for 5 min at 4°C, and the pellet was washed once with PBS. After centrifugation at 50 x g, the cells were resuspended in PBS and left on ice for 45 min. A second Percoll purification was then performed to further purify the cell suspension. The final pellet was resuspended in W10 plating medium. W10 medium is WEM supplemented with 10% FBS, 1% penicillin/streptomycin (cat#15140–12, Life Technologies), 1% 200 mM L-glutamine (cat#25030–081, Life Technologies), 0.1% Gentamicin reagent solution (50 mg/mL) (cat#15750–060, Life Technologies), and 0.1% Corning ITS premix (cat#354350, Corning). Viable cells were counted using trypan blue. Cell viability was typically above 95%, and minimal cell clumping was observed. When cell clumps were present, the suspension was filtered through a 40 μm cell strainer. For adoptive transplant into new FNRG recipient mice, cells were centrifuged at 50 x g for 5 min at 4°C and resuspended in cold PBS. For each mouse, 65 μL of cell suspension of 0.5 to 1 million cells was injected. For mpPHH culture and maintenance, cells were kept in Hepatocyte Culture Medium (HCM) (cat#CC3198, Lonza) supplemented with 2% DMSO (cat#4-x, ATCC), 1% gentamicin, and 0.5% ciprofloxacin (cat#17850-5G-F; Sigma-Aldrich). Media was typically replaced every 2-3 days. Distinct mpPHH mouse numbers throughout the study refer to independent hepatocyte preparations isolated from separately humanized FNRG mice engrafted with hepatocytes originating from the same human donor unless otherwise indicated.

### Primary human hepatocytes (PHH)

Cryopreserved (unpassaged) PHH (cat#HUM4129, Lonza) were first thawed at 37°C and then transferred to 50 mL W10. After centrifugation at 50 × g, the cells were resuspended in W10, counted, and seeded on collagen-coated plates (BD Biosciences) in W10 plating medium. The cells were equally distributed by shaking on a flat surface and then left on the bench at room temperature for 45 min. Once the cells settled and were evenly distributed, they were transferred to a humidified 37°C incubator. To avoid concentrating cells in the middle of the well, we seeded the cells with excess medium. The next day, the cells were washed once with WEM to remove any cell debris and serum. For maintenance medium, we used HCM supplemented with 2% DMSO, 1% gentamicin, and 0.5% ciprofloxacin. Unless otherwise stated, the medium was changed every other day.

### Enzyme-linked immunosorbent assay (ELISA) for human albumin

Serum was collected from the tail vein or retro-orbital plexus of humanized FNRG mice. Human albumin (hAlb) levels were quantified using a sandwich ELISA assay (Bethyl Laboratories) to determine the extent of human hepatocyte repopulation, as serum hAlb levels correlate with liver humanization in FNRG mice. Humanized FNRG mice with serum hAlb levels of approximately 10 mg/mL were considered to exhibit >90% liver repopulation and were preferentially used for mpPHH isolation. To assess hepatocyte health in mpPHH cultures, culture supernatants were collected at the indicated time points and hAlb levels were measured using the same ELISA assay. ELISA plates (Thermo Fisher Scientific, cat# 4424040) were coated with goat anti-human albumin polyclonal antibody (Bethyl Laboratories, cat# A80-129A), and bound hAlb was detected using HRP-conjugated detection antibody (Bethyl Laboratories, cat# A80-229P). Signal was developed using TMB substrate (Sigma Millipore, cat# T0440-100ML) and stopped with sulfuric acid (Fisher Chemical, cat# A300-500).

### HBeAg and HBsAg chemiluminescence assays (CLIA)

Secreted HBeAg and HBsAg were quantified using the HBeAg and HBsAg chemiluminescence immunoassays (CLIA). Kits were purchased from Diasino (cat# 0024082 for HBeAg and 0024031 for HBsAg). Assays were performed according to the manufacturer’s protocol.

### HBV infection and detection in mpPHH

Infection and detection were conducted as previously described^12,55,56^. HBV inoculum was generated from HepDE19 cells^57^, a tetracycline-regulated HBV-producing hepatoma cell line containing an integrated replication-competent HBV genome (genotype D). Viral stocks were produced from cultured cells rather than patient-derived samples. In summary, the supernatant of confluent HBV-producing HepDE19 cells was harvested every other day for three weeks and concentrated using filter devices (cat#C3043, MilliporeSigma). The concentrated stock was kept at -80°C. MpPHH were infected in HCM with 2% DMSO in the presence of 4% polyethylene glycol 8000 (cat#812868-250G, Sigma) and spinoculated for 1h at 1,000 x g at 37°C. Cells were washed after 24 h and, subsequently, the media was changed every other day. On day seven post-infection, the supernatant was collected, and HBsAg was analyzed using a chemiluminescence immunoassay (cat#0024031, Diasino) according to the manufacturer’s instructions. A FLUOstar Omega luminometer was used to read the plates. For *in vivo* infection, serum from humanized mice infected with genotype D from HepDE19-derived stock was injected intravenously into NTCP KO humanized mice.

### HBV infection and detection in HepG2-NTCP cells

HepG2-NTCP cells were maintained in Dulbecco’s Modified Eagle Medium (DMEM; Cytiva, SH30081.95) supplemented with 10% fetal bovine serum (FBS; Cytiva, SH30056.03), 1% non-essential amino acids (NEAA; Gibco, 11140050), and 2% DMSO at 37°C in a humidified 5% CO₂ incubator. For infection experiments, cells were seeded at 25,000 cells per well in collagen-coated 96-well plates in DMEM supplemented with 10% FBS and 1% NEAA. The following day, cultures were switched to DMEM supplemented with 3% FBS, 1% NEAA, and 2% DMSO. Twenty-four hours later, cells were infected with HBV derived from HepDE19 cells or a virus-free control inoculum in the presence of 4% polyethylene glycol 8000 (cat#812868-250G, Sigma) and 2% DMSO. Infection was enhanced by spinoculation at 1,000 x g for 1 h at 37°C, followed by incubation with the inoculum for an additional 24 h. The inoculum was then removed, cells were washed three times with PBS, and fresh culture medium containing 3% FBS, 1% NEAA, and 2% DMSO was added. Media were replenished 1 and 4 days post-infection. At 7 days post-infection, culture supernatants were collected, and cells were fixed in 4% paraformaldehyde (PFA) for downstream analyses.

### Cell viability assay

Cell viability was assessed using the resazurin-based PrestoBlue HS Cell Viability Reagent (cat#P50201, Thermo Fisher Scientific) according to the manufacturer’s instructions, including an empty control.

### Synthetic RNP delivery of CRISPR-Cas9

MpPHH were transfected with recombinant Cas9 protein (cat#CAS12207, Horizon Discovery) complexed with synthetic sgRNAs (Synthego). SgRNAs were resuspended in RNAse-free water. SgRNAs and recombinant Cas9 were thawed on ice and merged in OptiMEM medium at 37°C for 5 minutes for RNP formation. DharmaFECT Duo transfection reagent was added to HCM containing 2% DMSO, but lacking antibiotics, at 37°C. Then, both suspensions were mixed at a 1:1 ratio and, after incubation at room temperature for 20 min and washing the cells once with WEM at 37°C, were added directly to each well (for a 96-well plate: 50 µL/well). Final concentrations were 1% for DharmaFECT Duo, 60 nM for Cas9, and 100 nM for sgRNA. Sequentially, cells were centrifuged at 1,000 x g for 1 h at 37°C (spinoculation) to enhance uptake. Following transfection, cells were maintained in HCM containing 2% DMSO, with medium changes every 2–3 days. Non-targeting and lethal control crRNAs were obtained from Horizon Discovery (Dharmacon Edit-R platform; catalog numbers U-007501-01-20, U-006000-01-20, and U-002005-20). The exact guide RNA sequences are proprietary and were not disclosed by the manufacturer. These controls are bioinformatically designed to lack complementarity to the human genome (non-targeting) or to induce cell death through targeting of essential genomic elements (lethal control).

### Arrayed sgRNA library and screening design

The arrayed sgRNA library (Synthego) used for screening comprised 85 genes selected for their reported or predicted roles in innate immunity, hepatocyte biology, and host–virus interactions. Each gene was targeted using 4 independent sgRNAs per well to minimize guide-specific effects. Synthetic sgRNAs were obtained from Synthego and arrayed in a 96-well plate format, with each well containing sgRNAs targeting a single gene. Non-targeting and lethal control sgRNAs were included on each plate as negative and positive controls, respectively. A complete list of all sgRNAs and their sequences is provided in **Supplementary Table 1**.

### Lentivirus-vector–mediated delivery of Cas9 and sgRNAs

Lentiviral particles were produced in HEK293T cells cultured in T175 flasks. Prior to transfection, culture medium was replaced with DMEM supplemented with 10% FBS and 1% non-essential amino acids (NEAA). For each T175 flask, sgRNA or Cas9 transfer plasmids were co-transfected together with the second-generation packaging plasmid psPAX2 (5.5 μg; Addgene #12260) and envelope plasmid pMD2.G (2 μg; Addgene #12259) using FuGENE 4K transfection reagent (45 μL; Promega, cat# E5911) in Opti-MEM (682 μL). Transfection mixtures were incubated at room temperature for 30 min before addition to HEK293T cultures. Viral supernatants were harvested at 48 and 72 h post-transfection, clarified by centrifugation, filtered through 0.45 μm filters, aliquoted, and stored at −80°C until use. For CRISPR experiments, mpPHH were transduced with lentiviral expression vectors encoding Cas9 and gene-specific sgRNAs. This approach is distinct from the synthetic RNP lipofection method described above, as Cas9 and sgRNAs are expressed intracellularly following transduction rather than delivered as pre-assembled ribonucleoprotein complexes. MpPHH were transduced in HCM supplemented with 2% DMSO and 4 μg/mL polybrene (Sigma-Aldrich, cat# TR-1003-G) by spinoculation at 1,000 × g for one hour at 37°C. For *SLC10A1* (NTCP) targeting, cells were co-transduced with a lentiviral vector expressing four sgRNAs targeting *SLC10A1* and a second lentiviral vector expressing Cas9-RFP under a CMV promoter, enabling visual assessment of transduction efficiency. For *CYP3A4* targeting, cells were co-transduced with a lentiviral sgRNA vector and a Cas9-blasticidin construct driven by an EF1α promoter, enabling selection-based enrichment of Cas9-expressing cells. The use of these two Cas9 constructs provided complementary advantages: the Cas9-RFP system enabled rapid visualization of transduction efficiency, whereas the Cas9-blasticidin system enriched edited cell populations, which was particularly important for experiments requiring sustained Cas9 expression and for downstream applications such as transplantation into mice. Following transduction, cells were maintained in HCM supplemented with 2% DMSO, with media changes every 2–3 days. Cells were harvested 14 days post-transduction to allow sufficient time for Cas9 and sgRNA expression, genome editing, and turnover of pre-existing target protein. Protein lysates were prepared in RIPA buffer (Thermo Fisher Scientific, cat# 89900) and analyzed by western blotting for NTCP or CYP3A4 expression. Unless otherwise specified, mpPHH were maintained in HCM supplemented with 2% DMSO throughout all CRISPR editing, infection, and functional assays.

### CYP3A4 activity assay

Enzyme activity was determined using a cell-based P450-Glo CYP3A4 Assay and Screening System with luciferin-IPA (cat#V9002, Promega) in DMSO-free media according to the manufacturer’s instructions, including an empty control.

### Western blot analysis

Cells were washed with PBS, scraped, pelleted, and lysed with RIPA Lysis and Extraction Buffer (Thermo Fisher Scientific, cat# 89900), followed by separation by SDS-PAGE and transfer to nitrocellulose membrane. Membranes were incubated overnight in primary antibody (Cas9: cat#ab191468, abcam; CYP3A4: cat#MA517064, Fisher; SLC10A1: cat#PA525614, Thermo Fisher Scientific; human albumin HRP: cat#A80-229P, Thermo Fisher Scientific; beta-actin: cat#A5316, Sigma-Aldrich; beta-tubulin: cat#HRP-66240, Proteintech), and 0.1% milk diluted in TBS-T. TBS-T was used to wash for one hour before incubation with secondary antibodies from AlexaFluor (Thermo Fisher) of the corresponding species. Pictures were taken with an Azure imager and processed with ImageJ. For clarity, all the uncropped WB images are shown in **Supplementary Fig. 6**.

### Immunofluorescence staining

For immunofluorescence staining, cells were fixed and stained with a primary antibody against human CYP3A4 (cat# MA517064, Fisher) and an Alexa Fluor 594 (Thermo Fisher) secondary antibody or anti-HBV core antibody (Cell Marque; Cat# 216A-16-ASR) and same secondary. Nuclei were stained with Hoechst. A Nikon Eclipse TE300 fluorescent microscope was utilized for imaging, and ImageJ for processing.

### Quantification and statistical analysis

All experiments were performed with at least three biological replicates unless otherwise indicated. Biological replicates represent independent wells or independent hepatocyte preparations, whereas technical replicates correspond to repeated measurements within the same experimental unit. Data are presented as mean ± SD. Statistical analyses were performed using GraphPad Prism (version 10). Comparisons between two groups were conducted using unpaired two-tailed Student’s t-tests. For comparisons involving more than two groups, one-way ANOVA followed by Tukey’s multiple comparisons test was applied. For arrayed CRISPR screening experiments, statistical significance was assessed using one-way ANOVA with Benjamini–Krieger–Yekutieli correction for multiple comparisons, controlling the false discovery rate (FDR) at 0.05. The exact statistical tests and sample sizes (n) used for each experiment are indicated in the corresponding figure legends. A P value < 0.05 was considered statistically significant.

## Acknowledgments

This work was supported by NIH grants R21AI176944, R21AI176944-S1, R01AI181682, R56AI182395, DP1DK139804 and DP1DA066168 to E.M.; R01AA027327 and P01HL160472 to Y.J.; R01AI143295, R01AI150275 to C.R.; R01AI190067 to C.R., Y.J., and E.M.; DZIF MD fellowship (BMBF, Germany) to A.S.; C.F. is the Berger Foundation Fellow of the Damon Runyon Cancer Research Foundation (DRG-2440-21). This work was supported by Stavros Niarchos Foundation (SNF) as part of its grant to the SNF Institute for Global Infectious Disease Research at the Rockefeller University (to C.M.R.). We thank Julie Vercauteren for graphical assistance.

## Contributions

A.S., A.A., G.D., P.P., A.M., Ev.M., A.C., C.F., Y.Z., K.L., C.Q., C.Z., Y.J., and E.M. performed the experiments. A.S., A.A., Y.J., and E.M. analyzed data. A.S. and A.A. wrote the manuscript. A.G. provided reagents and assisted in methodology for sgRNAs; C.R., W.S., C.F., A.A., G.D., Ev.M., and Y.J. reviewed and edited the manuscript; E.M. designed the research, provided overall supervision, and edited the manuscript.

## References

1 Jacek, R. W., Anna, V., Agneta, N. & Per, A. In-depth quantitative analysis and comparison of the human hepatocyte and hepatoma cell line HepG2 proteomes. Journal of Proteomics 136, 234–247 (2016). 10.1016/j.jprot.2016.01.016

2 Kullak-Ublick, G. A. et al. Drug-induced liver injury: recent advances in diagnosis and risk assessment. Gut 66, 1154–1164 (2017). 10.1136/gutjnl-2016-313369

3 Ren, M. et al. The intersection of virus infection and liver disease: A comprehensive review of pathogenesis, diagnosis, and treatment. WIREs Mechanisms of Disease 16, e1640 (2024). 10.1002/wsbm.1640

4. World Health Organisation Hepatitis B Fact Sheet 2022 [Updated 23 July 2025] [(accessed on 24 November 2025)]*. Available online:* https://www.who.int/en/news-room/fact-sheets/detail/hepatitis-b.

5 Wilkening, S., Stahl, F. & Bader, A. COMPARISON OF PRIMARY HUMAN HEPATOCYTES AND HEPATOMA CELL LINE HEPG2 WITH REGARD TO THEIR BIOTRANSFORMATION PROPERTIES. Drug Metabolism and Disposition 31, 1035–1042 (2003). 10.1124/dmd.31.8.1035

6 Kaur, I. et al. Primary Hepatocyte Isolation and Cultures: Technical Aspects, Challenges and Advancements. LID - 10.3390/bioengineering10020131 [doi] LID - 131.

7 Azuma, H. et al. Robust expansion of human hepatocytes in Fah-/-/Rag2-/-/Il2rg-/- mice. Nat Biotechnol 25, 903–910 (2007). 10.1038/nbt1326

8. Bissig, K. D. et al. Human liver chimeric mice provide a model for hepatitis B and C virus infection and treatment.

9 Mercer, D. F. et al. Hepatitis C virus replication in mice with chimeric human livers. Nature Medicine 7, 927–933 (2001). 10.1038/90968

10. Dandri, M. et al. Repopulation of mouse liver with human hepatocytes and in vivo infection with hepatitis B virus.

11 de Jong, Y. P. Mice Engrafted with Human Liver Cells. Semin Liver Dis 44, 405–415 (2024). 10.1055/s-0044-1790601

12 Michailidis, E. et al. Expansion, in vivo-ex vivo cycling, and genetic manipulation of primary human hepatocytes. Proc Natl Acad Sci U S A 117, 1678–1688 (2020). 10.1073/pnas.1919035117

13 Bock, C. High-content CRISPR screening. Nature Reviews Methods Primers 2, 9 (2022). 10.1038/s43586-022-00098-7

14 Erard, N., Knott, S. R. V. & Hannon, G. J. A CRISPR Resource for Individual, Combinatorial, or Multiplexed Gene Knockout. Molecular Cell 67, 348–354.e344 (2017). 10.1016/j.molcel.2017.06.030

15 Metzakopian, E. et al. Enhancing the genome editing toolbox: genome wide CRISPR arrayed libraries. Scientific Reports 7, 2244 (2017). 10.1038/s41598-017-01766-5

16 Schmidt, T., Schmid-Burgk, J. L. & Hornung, V. Synthesis of an arrayed sgRNA library targeting the human genome. Scientific Reports 5, 14987 (2015). 10.1038/srep14987

17 Rathbone, T. et al. Electroporation-Mediated Delivery of Cas9 Ribonucleoproteins Results in High Levels of Gene Editing in Primary Hepatocytes. Crispr j 5, 397–409 (2022). 10.1089/crispr.2021.0134

18 Zabulica, M. et al. Correction of a urea cycle defect after ex vivo gene editing of human hepatocytes. Mol Ther 29, 1903–1917 (2021). 10.1016/j.ymthe.2021.01.024

19 Rouet, R. et al. Receptor-Mediated Delivery of CRISPR-Cas9 Endonuclease for Cell-Type-Specific Gene Editing. J Am Chem Soc 140, 6596–6603 (2018). 10.1021/jacs.8b01551

20 Tsukamoto, T. et al. Generation of the Adenovirus Vector-Mediated CRISPR/Cpf1 System and the Application for Primary Human Hepatocytes Prepared from Humanized Mice with Chimeric Liver. Biol Pharm Bull 41, 1089–1095 (2018). 10.1248/bpb.b18-00222

21 Mangeot, P. E. et al. Genome editing in primary cells and in vivo using viral-derived Nanoblades loaded with Cas9-sgRNA ribonucleoproteins.

22 Chong, Z. X., Yeap, S. K. & Ho, W. Y. Transfection types, methods and strategies: a technical review. PeerJ 9, e11165 (2021). 10.7717/peerj.11165

23 Verma, R. S., Giannola, D., Shlomchik, W. & Emerson, S. G. Increased efficiency of liposome-mediated transfection by volume reduction and centrifugation. Biotechniques 25, 46–49 (1998). 10.2144/98251bm09

24 Davidson, M. D. & Khetani, S. R. Intermittent Starvation Extends the Functional Lifetime of Primary Human Hepatocyte Cultures. Toxicol Sci 174, 266–277 (2020). 10.1093/toxsci/kfaa003

25 Neuhaus, B. et al. Nanoparticles as transfection agents: a comprehensive study with ten different cell lines. Rsc Advances 6, 18102–18112 (2016).

26. Buyl, K., De Kock, J., Bolleyn, J., Rogiers, V., Vanhaecke, T. Measurement of Albumin Secretion as Functionality Test in Primary Hepatocyte Cultures, bookTitle= Protocols in In Vitro Hepatocyte Research. (Springer New York, 2015).

27. Julia, R., Birgit, M. W., Espen, M. & Magnus, I.-S.

28 Peng, Z. et al. Requirments for primary human hepatocyte. Cell Prolif 55, e13147 (2022). 10.1111/cpr.13147

29 Shimada, T., Yamazaki, H., Mimura, M., Inui, Y. & Guengerich, F. P. Interindividual variations in human liver cytochrome P-450 enzymes involved in the oxidation of drugs, carcinogens and toxic chemicals: studies with liver microsomes of 30 Japanese and 30 Caucasians. J Pharmacol Exp Ther 270, 414–423 (1994).

30 Peter Guengerich, F. Enzymatic Oxidation of Xenobiotic Chemical. Critical Reviews in Biochemistry and Molecular Biology 25, 97–153 (1990). 10.3109/10409239009090607

31 Gonzalez, F. & Nebert, D. Evolution of the P450 gene superfamily: animal-plant ‘warfare’, molecular drive and human genetic differences in drug oxidation. Trends Genet. 6, 182–186 (1990). 10.1016/0168-9525(90)90174-5

32 Zanger, U. M., Turpeinen, M., Klein, K. & Schwab, M. Functional pharmacogenetics/genomics of human cytochromes P450 involved in drug biotransformation. Analytical and Bioanalytical Chemistry 392, 1093–1108 (2008). 10.1007/s00216-008-2291-6

33 Mancio-Silva, L. et al. Improving Drug Discovery by Nucleic Acid Delivery in Engineered Human Microlivers. Cell Metab 29, 727–735.e723 (2019). 10.1016/j.cmet.2019.02.003

34 Isom, H. C., Secott, T., Georgoff, I., Woodworth, C. & Mummaw, J. Maintenance of differentiated rat hepatocytes in primary culture. Proc Natl Acad Sci U S A 82, 3252– 3256 (1985). 10.1073/pnas.82.10.3252

35 Gripon, P. et al. Hepatitis B virus infection of adult human hepatocytes cultured in the presence of dimethyl sulfoxide. J Virol 62, 4136–4143 (1988). 10.1128/jvi.62.11.4136-4143.1988

36 Kamdem, L. K., Meineke, I., Gödtel-Armbrust, U., Brockmöller, J. & Wojnowski, L. Dominant Contribution of P450 3A4 to the Hepatic Carcinogenic Activation of Aflatoxin B1. Chemical Research in Toxicology 19, 577–586 (2006). 10.1021/tx050358e

37 Ishida, Y. et al. Detection of acute toxicity of aflatoxin B1 to human hepatocytes in vitro and in vivo using chimeric mice with humanized livers. PLoS One 15, e0239540 (2020). 10.1371/journal.pone.0239540

38 de Jong, Y. P. et al. Broadly neutralizing antibodies abrogate established hepatitis C virus infection. Sci Transl Med 6, 254ra129 (2014). 10.1126/scitranslmed.3009512

39 Braun, R. J. & Zischka, H. Mechanisms of Cdc48/VCP-mediated cell death: from yeast apoptosis to human disease. Biochim Biophys Acta 1783, 1418–1435 (2008). 10.1016/j.bbamcr.2008.01.015

40 Hyrina, A. et al. A Genome-wide CRISPR Screen Identifies ZCCHC14 as a Host Factor Required for Hepatitis B Surface Antigen Production. Cell Rep 29, 2970–2978 e2976 (2019). 10.1016/j.celrep.2019.10.113

41 Yan, H. et al. Sodium taurocholate cotransporting polypeptide is a functional receptor for human hepatitis B and D virus. Elife 1, e00049 (2012). 10.7554/eLife.00049

42 McCaughey, J. et al. A general role for TANGO1, encoded by MIA3, in secretory pathway organization and function. J Cell Sci 134 (2021). 10.1242/jcs.259075

43 Verrier, E. R. et al. Hepatitis B Virus Evasion From Cyclic Guanosine Monophosphate-Adenosine Monophosphate Synthase Sensing in Human Hepatocytes. Hepatology 68, 1695–1709 (2018). 10.1002/hep.30054

44 Sato, S. et al. The RNA sensor RIG-I dually functions as an innate sensor and direct antiviral factor for hepatitis B virus. Immunity 42, 123–132 (2015). 10.1016/j.immuni.2014.12.016

45 Guo, H. et al. Activation of pattern recognition receptor-mediated innate immunity inhibits the replication of hepatitis B virus in human hepatocyte-derived cells. J Virol 83, 847–858 (2009). 10.1128/jvi.02008-08

46 Liu, Y. et al. Interferon-inducible ribonuclease ISG20 inhibits hepatitis B virus replication through directly binding to the epsilon stem-loop structure of viral RNA. PLoS Pathog 13, e1006296 (2017). 10.1371/journal.ppat.1006296

47 Benjamini, Y., Krieger, A. M. & Yekutieli, D. Adaptive linear step-up procedures that control the false discovery rate. Biometrika 93, 491–507 (2006). 10.1093/biomet/93.3.491

48 Kabbani, M. et al. Human hepatocyte PNPLA3-148M exacerbates rapid non-alcoholic fatty liver disease development in chimeric mice. Cell Rep 40, 111321 (2022). 10.1016/j.celrep.2022.111321

49 Yin, J.-A. et al. Arrayed CRISPR libraries for the genome-wide activation, deletion and silencing of human protein-coding genes. Nature Biomedical Engineering 9, 127– 148 (2025). 10.1038/s41551-024-01278-4

50 Haapaniemi, E., Botla, S., Persson, J., Schmierer, B. & Taipale, J. CRISPR-Cas9 genome editing induces a p53-mediated DNA damage response. Nat Med 24, 927–930 (2018). 10.1038/s41591-018-0049-z

51 Yu, J., Liang, C. & Liu, S. L. Interferon-inducible LY6E Protein Promotes HIV-1 Infection. J Biol Chem 292, 4674–4685 (2017). 10.1074/jbc.M116.755819

52 Hackett, B. A. & Cherry, S. Flavivirus internalization is regulated by a size-dependent endocytic pathway. Proc Natl Acad Sci U S A 115, 4246–4251 (2018). 10.1073/pnas.1720032115

53 Mar, K. B. et al. LY6E mediates an evolutionarily conserved enhancement of virus infection by targeting a late entry step. Nat Commun 9, 3603 (2018). 10.1038/s41467-018-06000-y

54 Pfaender, S. et al. LY6E impairs coronavirus fusion and confers immune control of viral disease. Nat Microbiol 5, 1330–1339 (2020). 10.1038/s41564-020-0769-y

55 Michailidis, E. et al. A robust cell culture system supporting the complete life cycle of hepatitis B virus. Sci Rep 7, 16616 (2017). 10.1038/s41598-017-16882-5

56 Jones, C. E. et al. Long-term 3D cell culture models for hepatitis B virus studies. Virology 600, 110265 (2024). 10.1016/j.virol.2024.110265

57 Cai, D. et al. Identification of disubstituted sulfonamide compounds as specific inhibitors of hepatitis B virus covalently closed circular DNA formation. Antimicrob Agents Chemother 56, 4277–4288 (2012). 10.1128/AAC.00473-12

