## Supplemental Figures for "A CRISPR-Cas9 platform for primary human hepatocytes enables arrayed screening and *in vivo* validation of HBV host factors"

^1^Laboratory of Virology and Infectious Disease, The Rockefeller University, New York, NY 10065, USA; ^2^Department of Infectious Diseases, Molecular Virology, Heidelberg University, Heidelberg, Germany; ^3^Department of Infectious Diseases, Respiratory Medicine and Critical Care, Charité - Universitätsmedizin Berlin, Corporate Member of Freie Universität Berlin and Humboldt-Universität zu Berlin (current address); ^4^Laboratory of Biochemical Pharmacology, Department of Pediatrics, Emory University, Atlanta, GA 30322, USA; ^5^Department of Gastroenterology, Hepatology and Infectious Diseases, University Hospital Dusseldorf, Medical Faculty at Heinrich Heine University Dusseldorf, Dusseldorf, Germany (current address); ^6^Division of Gastroenterology and Hepatology, Weill Cornell Medicine, New York, NY 10065, USA; ^7^Institute of Neuropathology, University of Zurich, Zurich, Switzerland; ^8^These authors contributed equally: Ansgar F. Stenzel and Antonis Athanasiadis; ^9^Corresponding author:

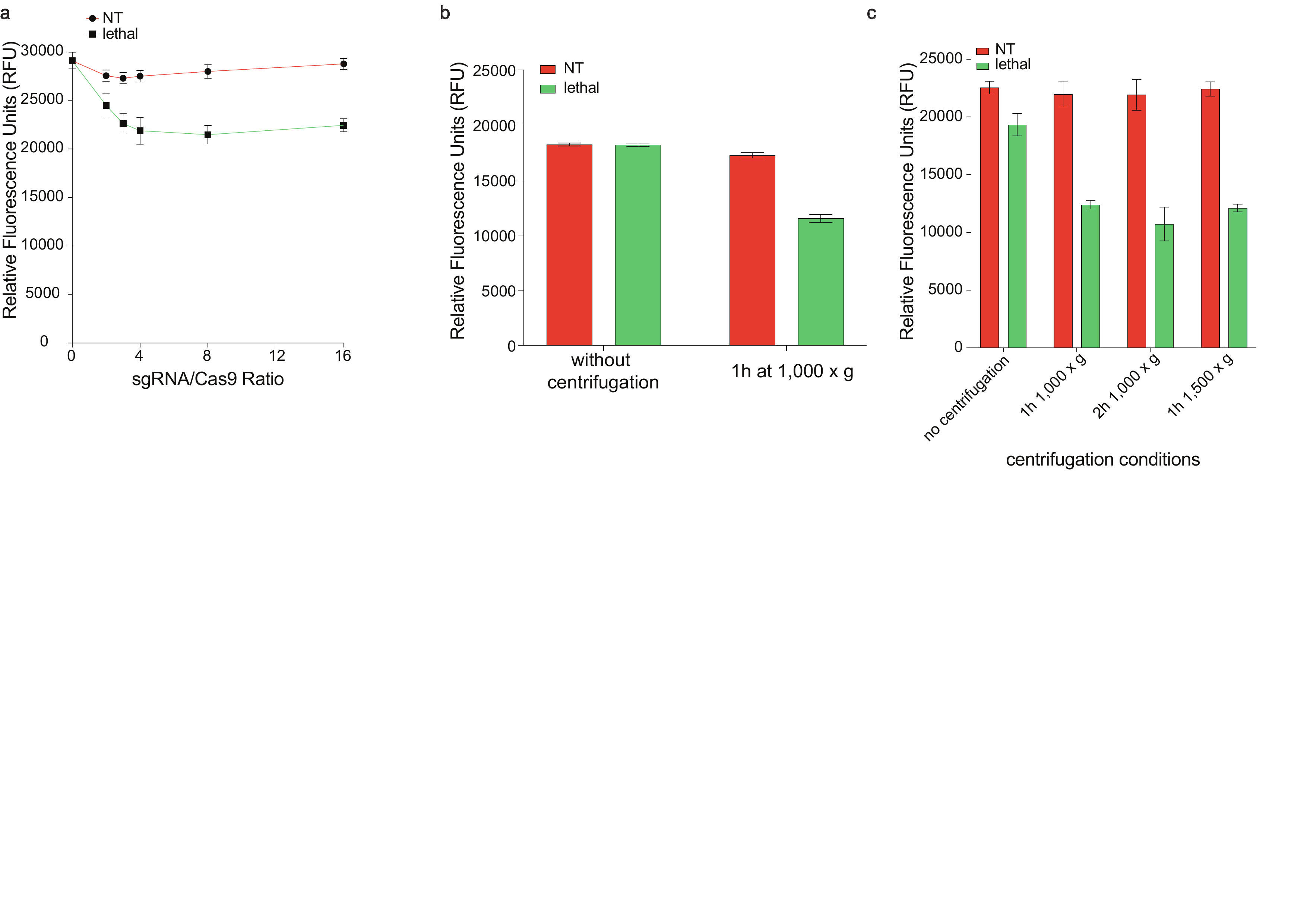

**Supplementary Fig. 1 | Optimization of lipofection-based RNP delivery in mpPHH. a** Optimization of sgRNA:Cas9 molar ratio using NT and lethal sgRNAs. Cell viability was measured by a fluorescence-based viability assay (PrestoBlue) and is shown as relative fluorescence units. **b** Effect of centrifugation-assisted delivery on RNP transfection outcome. mpPHH were transfected with NT or lethal sgRNAs with or without centrifugation at 1,000 × g for 1 h, and viability was measured by relative fluorescence units. **c** Optimization of centrifugation conditions for RNP delivery. mpPHH were transfected with NT or lethal sgRNAs and subjected to the indicated centrifugation conditions. Cell viability was measured by relative fluorescence units. Data are shown as mean ± SD.

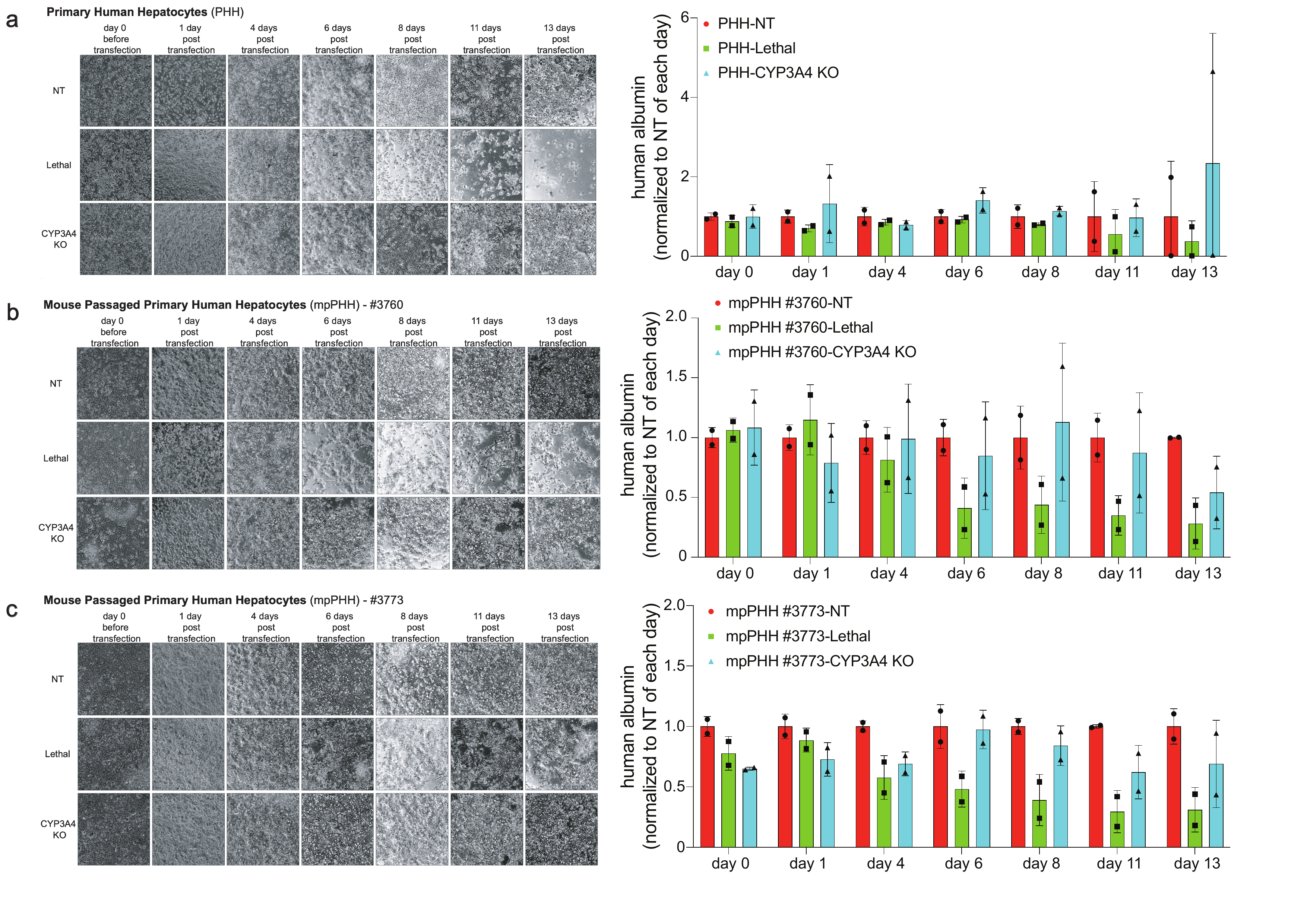

**Supplementary Fig. 2 | Longitudinal morphology and albumin secretion after RNP editing in PHH and mpPHH. a** Representative bright-field images and longitudinal human albumin measurements from unpassaged PHH transfected with NT, lethal, or *CYP3A4*-targeting sgRNAs. Images were acquired at the indicated time points after RNP lipofection, and hAlb secretion was normalized to the NT condition at each corresponding time point. **b** Representative bright-field images and longitudinal hAlb measurements from mpPHH donor #3760 transfected with NT, lethal, or *CYP3A4*-targeting sgRNAs. **c** Representative bright-field images and longitudinal hAlb measurements from mpPHH donor #3773 transfected with NT, lethal, or *CYP3A4*-targeting sgRNAs. Across PHH and mpPHH cultures, NT and *CYP3A4*-targeted cells maintained hepatocyte morphology and albumin secretion over time, whereas lethal sgRNA-transfected cells showed progressive loss of viability. Data are shown as mean ± SD.

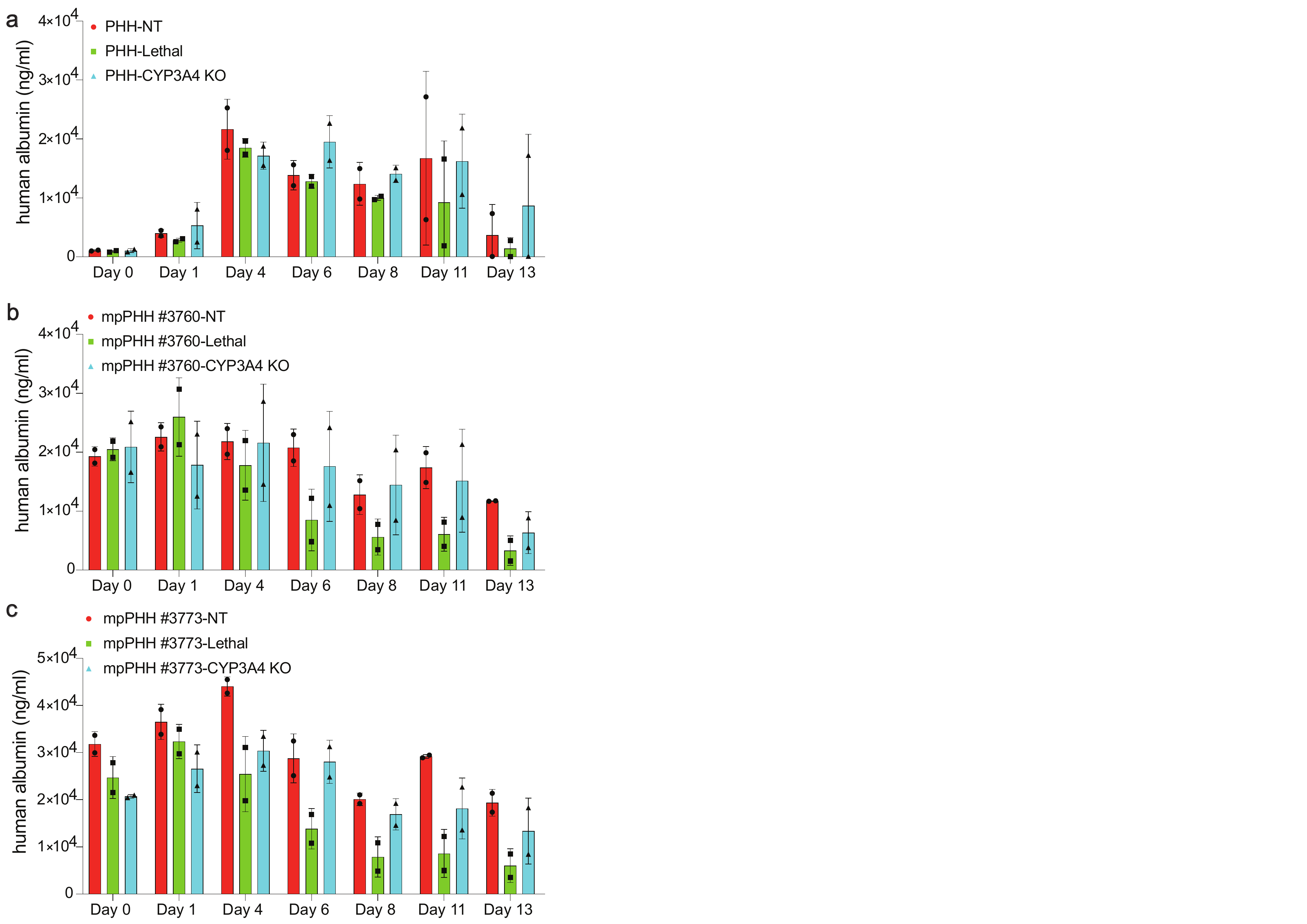

**Supplementary Fig. 3 | Absolute human albumin secretion after RNP editing in PHH and mpPHH. a** Absolute human albumin (hAlb) concentrations (ng/ml) from unpassaged PHH transfected with NT, lethal, or *CYP3A4*-targeting sgRNAs, measured longitudinally at the indicated time points after RNP lipofection and shown here without normalization to the NT condition (normalized values for the same experiment are shown in Supplementary Fig. 2a). **b** Absolute hAlb concentrations from mpPHH donor #3760 transfected with NT, lethal, or *CYP3A4*-targeting sgRNAs (normalized values shown in Supplementary Fig. 2b). **c** Absolute hAlb concentrations from mpPHH donor #3773 transfected with NT, lethal, or *CYP3A4*-targeting sgRNAs (normalized values shown in Supplementary Fig. 2c). Data are shown as mean ± SD.

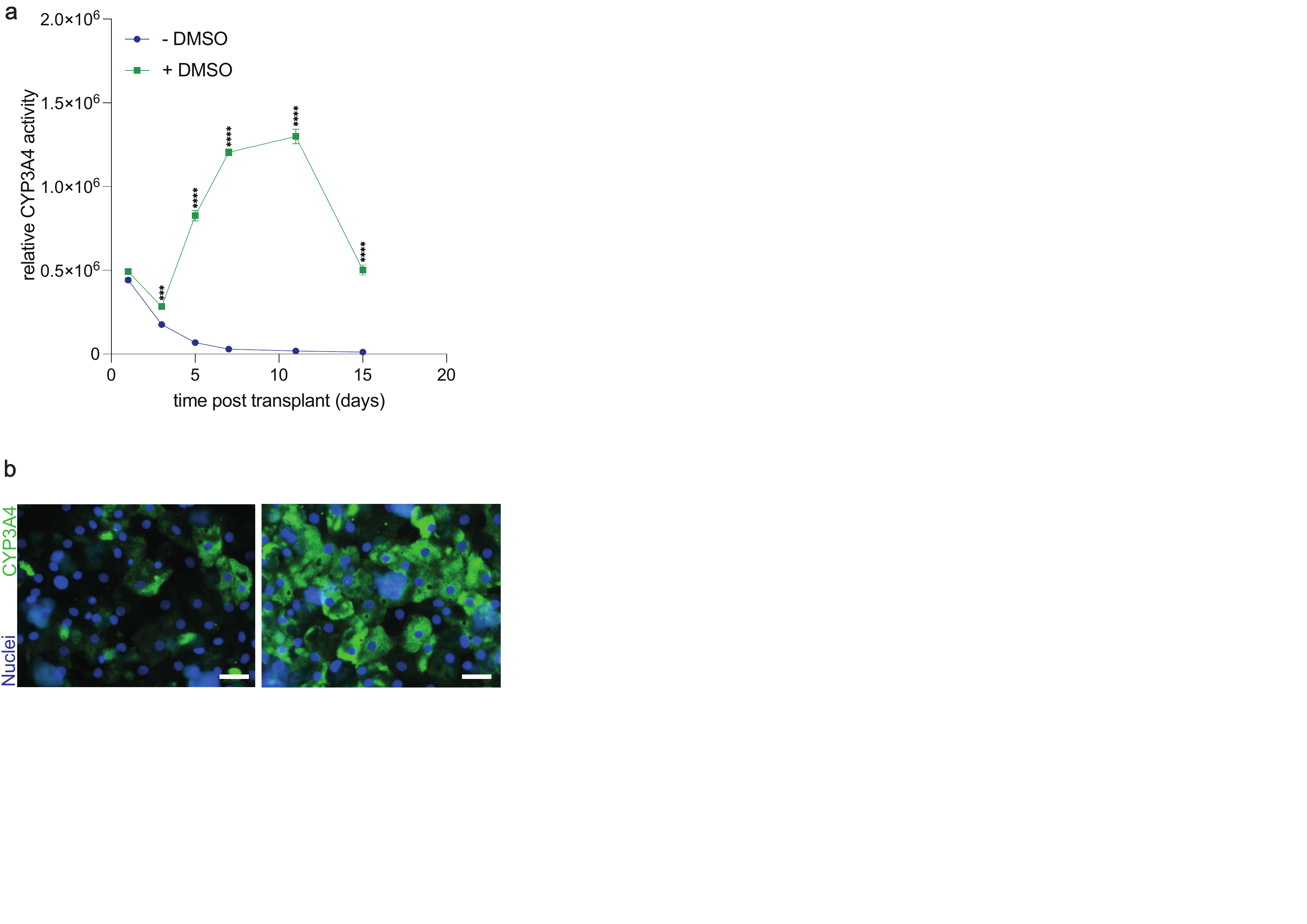

**Supplementary Fig. 4 | DMSO preserves *CYP3A4* activity and expression in cultured mpPHH. a** Longitudinal *CYP3A4* enzymatic activity in mpPHH cultured with or without 2% DMSO. Activity was measured using the luciferin-IPA substrate and is shown as relative activity over time. **b** Representative immunofluorescence images of mpPHH cultured with or without 2% DMSO and stained for CYP3A4 protein and nuclei. Scale bars, 50 μm. Data are shown as mean ± SD. Statistical significance was assessed as indicated in Methods. *P < 0.05, ***P < 0.001, ****P < 0.0001.

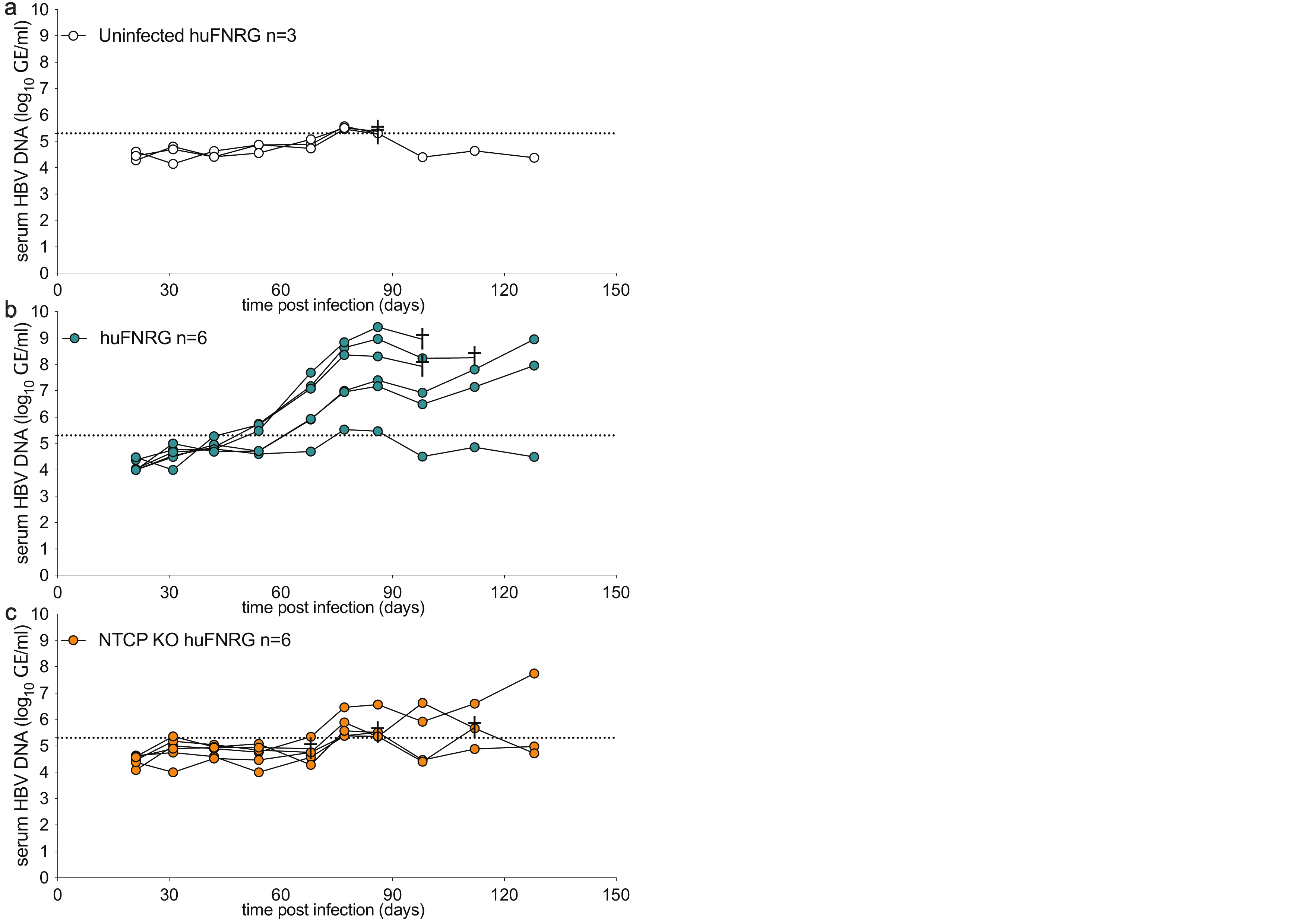

**Supplementary Fig. 5 | Individual serum HBV DNA trajectories underlying Fig. 5c. a** Serum HBV DNA levels in uninfected huFNRG mice (n=3); mice 4059 and 4066 were last measured at day 86. **b** Serum HBV DNA levels in HBV-infected huFNRG mice (n=6); mice 4055 and 4067 were last measured at day 98 and mouse 4069 at day 112. **c** Serum HBV DNA levels in NTCP-KO HBV-infected huFNRG mice (n=6); mice 3404, 3411, and 4350 were last measured at days 68, 86, and 112, respectively. Data are the same underlying values shown aggregated in Fig. 5c, plotted here by individual animal for clarity. The dotted line indicates the assay limit of quantification. † marks the final recorded time point for an animal that did not complete the full 130-day observation period: in **b**, marked animals were euthanized for planned liver harvest after reaching peak viremia to generate HBV-infected mpPHH; in **a** and **c**, marked animals either died spontaneously or were euthanized per humane endpoint criteria.

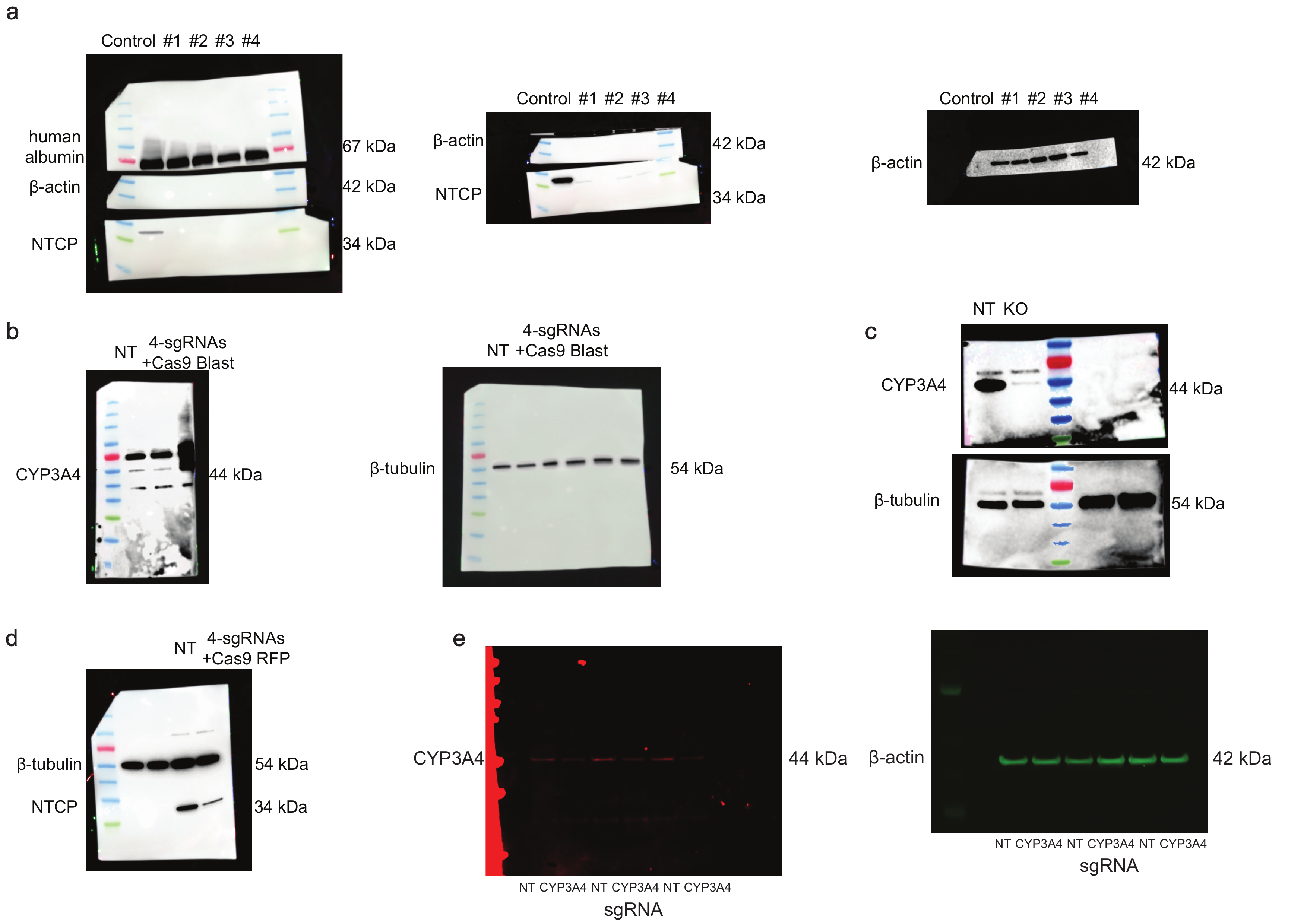

**Supplementary Fig. 6 | Uncropped western blots corresponding to main figures.** Blots include analyses of NTCP, CYP3A4 protein, human albumin, β-actin, and β-tubulin expression from the indicated experiments. Molecular weight markers and full membrane regions are shown where available. **a** is the western blot from Fig. 5, **b** from Fig. 6, **c** from Fig. 1, **d** from Fig. 6, and **e** from Fig. 2.
